# Development of Multiomics *in situ* Pairwise Sequencing (MiP-Seq) for Single-cell Resolution Multidimensional Spatial Omics

**DOI:** 10.1101/2023.01.07.523058

**Authors:** Xiaofeng Wu, Weize Xu, Lulu Deng, Yue Li, Zhongchao Wang, Leqiang Sun, Anran Gao, Haoqi Wang, Xiaodan Yang, Chengchao Wu, Yanyan Zou, Keji Yan, Zhixiang Liu, Lingkai Zhang, Guohua Du, Liyao Yang, Da Lin, Ping Wang, Yunyun Han, Zhenfang Fu, Jinxia Dai, Gang Cao

## Abstract

Delineating the spatial multiomics landscape will pave the way to understanding the molecular basis of physiology and pathology. However, current spatial omics technology development is still in its infancy. Here, we developed a high-throughput multiomics *in situ* pairwise sequencing (MiP-Seq) strategy to efficiently decipher multiplexed DNAs, RNAs, proteins, and small biomolecules at subcellular resolution. We delineated dynamic spatial gene profiles in the hypothalamus using MiP-Seq. Moreover, MiP-Seq was unitized to detect tumor gene mutations and allele-specific expression of parental genes and to differentiate sites with and without the m6A RNA modification at specific sites. MiP-Seq was combined with *in vivo* Ca^2+^ imaging and Raman imaging to obtain a spatial multiomics atlas correlated to neuronal activity and cellular biochemical fingerprints. Importantly, we proposed a “signal dilution strategy” to resolve the crowded signals that challenge the applicability of *in situ* sequencing. Together, our method improves spatial multiomics and precision diagnostics and facilitates analyses of cell function in connection with gene profiles.

---

Multicellular organisms elegantly orchestrate the cotranscription of the appropriate genes in specific cells ^1-3^. These spatial transcription profiles govern the functions of distinct cells to complete sophisticated physiological tasks. Delineation of comprehensive spatial omics will pave the way to understanding the molecular function of specific cells and greatly advance precision diagnostics by, for example, characterizing the immune microenvironment before cancer therapy ^4-8^. Recently, state-of-the-art spatial transcriptomics methods have been developed for highly multiplexed RNA *in situ* detection: these methods include SeqFISH ^9-11^, MERFISH ^12, 13^, ISS ^14-18^, FISSEQ ^19^, STARmap ^20^, INSTA-seq ^21^, ExSeq ^22, 23^, Slide-seq ^24, 25^, HDST ^26^, and DBiT-seq ^27^. However, spatial-omics technologies are still in their infancy. For example, most methods can be used to decipher only the spatial information of one kind of biomolecule, many rounds of sequencing are required for high-throughput spatial omics. Methods to improve the capture efficiency, throughput capacity, cellular resolution, and optical crowding and to reduce the cost of these experiments are desperately needed ^28-30^. In this study, we developed a high-throughput multiomics *in situ* pairwise sequencing (MiP-Seq) method to efficiently detect multiplexed DNAs, RNAs, proteins and biomolecules in brain tissue at subcellular resolution.

The *in situ* detection of DNA and RNA by MiP-Seq is realized with direct padlock probes that target nucleic acids, and the detection of proteins and biomolecules is realized with padlock probes that detect antibody-conjugated nucleic acids (Fig. 1A). First, we applied MiP-Seq to spatial RNA profiling, and the process involved padlock probe hybridization, *in situ* rolling-circle amplification (RCA) and dual barcode sequencing via ligation (Fig. 1B). Notably, each padlock probe contains two barcodes (B1 and B2), which increases its signal-decoding capacity. Considering previous studies^18, 31-34^, we compared different ligases (Fig. S1) and selected SplintR to ligate RCA templates, which enhanced RCA template ligation efficiency. This method of ligation depends on precisely matched base pairing, which allows single-nucleotide polymorphisms (SNP) in target RNA to be detected by MiP-Seq. Owing to the target-dependent ligation of the padlock probe and initiator primer (IP) complementarity to both the target RNA and padlock probe, RCA was firmly performed *in situ* together with the target (Fig. 1B). For high-throughput detection, the *in situ* rolling amplified products (RCPs) were subjected to multiple rounds of sequencing via ligation and signal decoding (Fig. 1B).

**Figure 1.**
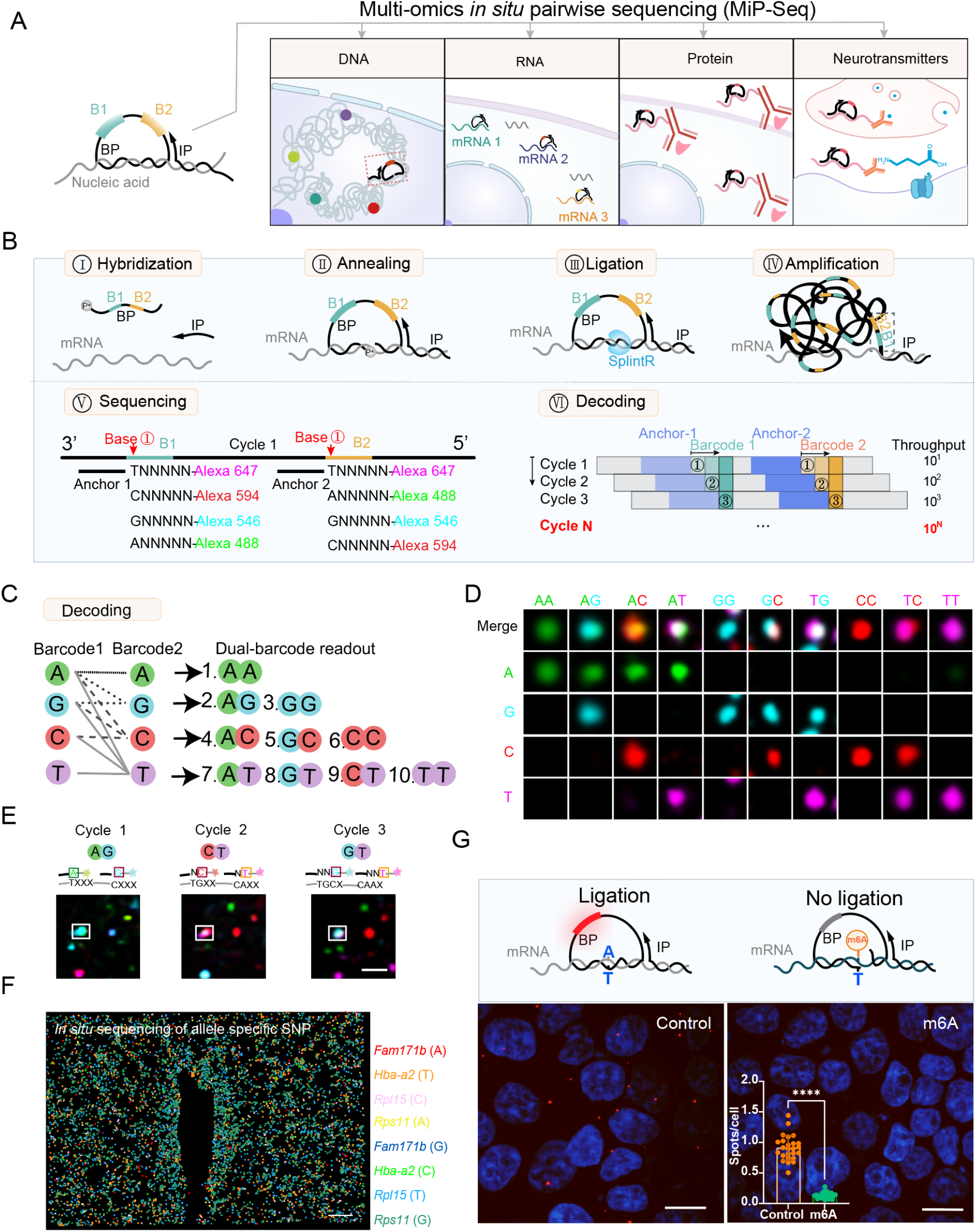
Application of multi-omics *in situ* pairwise sequencing (MiP-Seq) for spatial biomolecules profile. (**A**) Diagram for the application of multi-omics *in situ* pairwise sequencing (MiP-Seq) in spatial omics profiling. (**B**) Flow diagram of MiP-Seq for RNA *in situ* detection. I. The dual-barcode primer (BP) and initiator primer (IP) are hybridized to the target RNA. II. BP and IP are annealed to the target mRNA to form a padlock structure. III. BP is circulized by RNA dependent DNA ligase SplintR to form rolling-circle amplification (RCA) template. IV. RCA of circled BP forms rolling-circle products. V. The dual-anchor primers are hybridized next to the dual-barcode sequence. Subsequently, the barcode sequences are interrogated by fluorescence labeled query probes through sequencing by ligation. VI. Ten dual-barcode readout are decoded in each round of sequencing, and 10^N^ codes can be interrogated after N rounds. (**C**) Illustration of ten dual-barcodes interrogation according to merged florescence signals from paired bases for each round sequencing. (**D**) Merged florescence signals for each of ten dual-barcodes. (**E**) Example of dual-barcode sequence interrogation. The white boxed dot showed the corresponding barcode reading for each of the three rounds. (**F**) *In situ* sequencing for allele-specific gene expression of *Fam171b, Hba-a2, Rpl15*, and *Rps11* in the paraventricular nucleus (PVN) of hypothalamus of offspring mice from C57BL/6 male mice crossed with BALB/c female mice. (**G**) The m6A methylated RNA and the unmethylated RNA were transfected into HEK293T cells, which can be differentiated by MiP-Seq. Statistical analysis of unmethylated and methylated RNA from 25 imaging fields (****P < 0.0001). Scale bars, 2 µm in E, 50 µm in F, 25 µm in G.

To increase the signal-decoding throughput capacity, we simultaneously sequenced the dual barcode base in each round. In this strategy, the combination of two of four fluorescence signals enables barcoding of ten different combinations within each cycle (Fig. 1B-1D). Thus, 10^N^ genes can be decoded via N rounds of sequencing (Fig. 1B), which dramatically increases the gene detection throughput in considerably fewer rounds of sequencing compared to that of other *in situ* sequencing methods (in which usually 4^N^ genes can be decoded) ^14, 19, 20, 22^. Owing to RCA, the signals of RCPs, represented as dots, can be clearly recognized and interpreted based on dual barcode sequencing (Fig. 1D-1E). To perform dual barcode sequencing, we tried two strategies, paired-end sequencing and pairwise sequencing (Fig. S2A-S2B). In the paired-end sequencing strategy, a dual barcode base was sequenced by one-anchor paired-end ligation sequencing (Fig. S2A). Co-occurrence and correlation were analyzed to verify the percentage of overlapping dual barcodes in HBMECs (Fig. S2C-S2E). The intensities of the dual-barcode fluorescence signals were found to be highly correlated, suggesting the reliability of the dual barcode sequencing method (Fig. S2D). These data were validated via cooccurrence analysis of dual fluorescence signals (Fig. S2E).

To obtain maximal dual base-calling accuracy, we performed a pairwise sequencing to decipher the dual barcode and compared the efficiency of this method with that of the paired-end sequencing strategy (Fig. S2A-S2B). As shown in Fig. S2F-S2H, pairwise sequencing led to a greater overlapping than the paired-end sequencing strategy. These data indicate that our pairwise sequencing-based MiP-Seq method led to more accurate dual base-calling, on top of high barcoding capacity. With the rapid evolution of highly sensitive microscopy techniques and deep learning-assisted imaging analysis, we believe the accuracy of dual base-calling can be further improved.

Next, we tested the efficiency of MiP-Seq in RNA detection by comparing the degree of human *GAPDH* and *mTOR* gene transcript detection obtained with MiP-Seq and with HCR-FISH 3.0 ^35^ (FISH with HCR amplification) (Fig. S3). First, for MiP-Seq, 2 target probes against *GAPDH* and *MTOR* transcripts were used, and the average detection efficiency per cell reached approximately 40% of that achieved with HCR3.0-FISH. By increasing the number of MiP-Seq targeting probes to 8 probes, the efficiency of *GAPDH* and *MTOR* gene transcript detection reached 96% and 56%, respectively. Further, the expression pattern of different genes in the mouse brain that was detected by MiP-Seq was similar to the pattern revealed through traditional *in situ* hybridization according to the Allen Institute brain map (Fig. S4A-S4B). Moreover, MiP-Seq enabled detection of gene transcripts at subcellular resolution, as indicated by the results showing the *Kif5a* mRNA signal in the cytoplasm and the *Snhg11* mRNA signal in the nucleus (Fig. S4C).

MiP-Seq firmly depends on the adjacent ligation to a precisely matched base pair with the target sequence to form the RCA template. Thus, it may be further utilized to detect targets with single base variance and carrying RNA modifications. As shown in Fig. S5A, we performed MiP-Seq to distinguish human and mouse β-actin mRNA with single nucleotide variation in cocultured HBMEC cells (human origin) with Bend3 cells (mouse origin). Moreover, we leveraged MiP-Seq to analyze multiplex allele-specific gene expression patterns in mouse brains. The SNPs of parental genes (*Fam171b, Rpsl1, Hba-a2*, and *Rpl15*) were first identified by RNA-seq, and then, MiP-Seq was applied to samples of mouse hippocampus and paraventricular nucleus of the hypothalamus (PVN). Uneven allele-specific gene expression patterns of parental genes were consistently detected; for example, the G allele of the *Rps11* gene and the T allele of the *Rpl15* and *Hba-a2* genes, which were preferentially expressed in both the hippocampus and PVN, were detected by MiP-Seq (Fig. 1F and Fig. S5B-S5D). These data may imply that the maternal and paternal chromosomes in the same cell might be undergoing different gene transcription regulation process.

As RNA modification may disrupt RCA template ligation, we hypothesized that MiP-Seq can be used to differentiate methylated RNA from unmodified RNA. To test this hypothesis, synthesized m6A-modified RNA and control RNA without modification were transfected into HEK293T cells and subjected to MiP-Seq. As shown in Fig. 1G, we observed positive signals for unmodified RNA, whereas we detected a negligible signal for m6A-modified RNA due to failed adjacent ligation at the methylated nucleotide site. Thus, MiP-Seq can be utilized to differentiate m6A methylated RNA at specific sites. Furthermore, we used MiP-Seq to detect wild-type (WT) and point mutations of multiple genes in tumors simultaneously (Fig. S5E-S5F). Based on our RNA-seq data, we applied MiP-Seq to WT and mutated *Eno1, Nptxr, Spp1, Tnc*, and *Tubala* genes in rat brains with C6 glioma (Fig. S5E). The expression of the mutant and WT genes was simultaneously detected in these glioma tissues through 10 barcode readouts in one round of *in situ* sequencing (Fig. S5F).

Next, we tried to employ MiP-Seq to realize multiple genes detection for large imaging field of view. As shown in Fig. 2A, the spatial transcriptomic landscape of ten genes (*Calb1, Gad1, Plp1, Neurod1, Cck, Lamp5, Ndnf, Pax2, Pcp4*, and *Penk*) in the whole sagittal section of mice brain was delineated by one round of MiP-Seq. Importantly, the expression patten of these genes detected by MiP-Seq is consistent with the results from Allen Institute of Brain Science (Fig. 2B and Fig. S6A-S6B). To further construct 3D gene expression architecture, 34 sagittal sections from the whole cerebellum with 200 µm interval were subjected to multiplexed genes detection by MiP-Seq (Fig. 2C-2D and movie S1). The reconstructed image showed the 3D view of the expression and distribution of *Calb1, Gad1, Neurod1*, and *Plp1* genes in the whole cerebellum (Fig. 2E and Fig. S6C). Moreover, we also optimized MiP-Seq for multiplex gene detection in whole mount tissue. As shown in Fig.2F-2G, the *in situ* gene expression pattern of *foxg1a, hoxb3* and *gad1b* genes were detected in the whole mount of zebrafish telencephalon. Owing to the high-throughput capacity of MiP-Seq, we mapped the spatial transcriptome of 100 genes in the nucleus of the horizontal limb of the diagonal band (HDB) in the hypothalamus by 2 rounds of sequencing (Fig. 3A). High-throughput image processing involves image registration, candidate spot calling, decoding, cell segmentation, and gene assignment. The spatial transcriptome analysis pipeline for high-throughput pairwise dual-color signal decoding is shown in Fig. S7. Our data revealed the spatial profiling of 100 genes in the HDB, including neurotransmitter-related genes, neuronal marker genes, and neuronal activity marker genes, at single-cell resolution (Fig. 3A and Fig. S8A-S8B). Based on 100 gene profile atlases, the unique spatial expression patterns of peptide hormone genes, neurotransmitter-related genes, and their corresponding receptor genes were identified, as shown in Fig. 3B and Fig. S8C. Through spatial gene coexpression analysis, we found that certain genes were spatially coexpressed (Fig. 3C). For example, a high spatial correlation was observed for *Snap25* and *Scg2* expression (Fig. 3D-3E). Interestingly, we also identified that some genes with certain spatially exclusive expression patterns, such as *Hap1* and *Pdgfra* (Fig. S8D-S8E).

**Fig 2.**
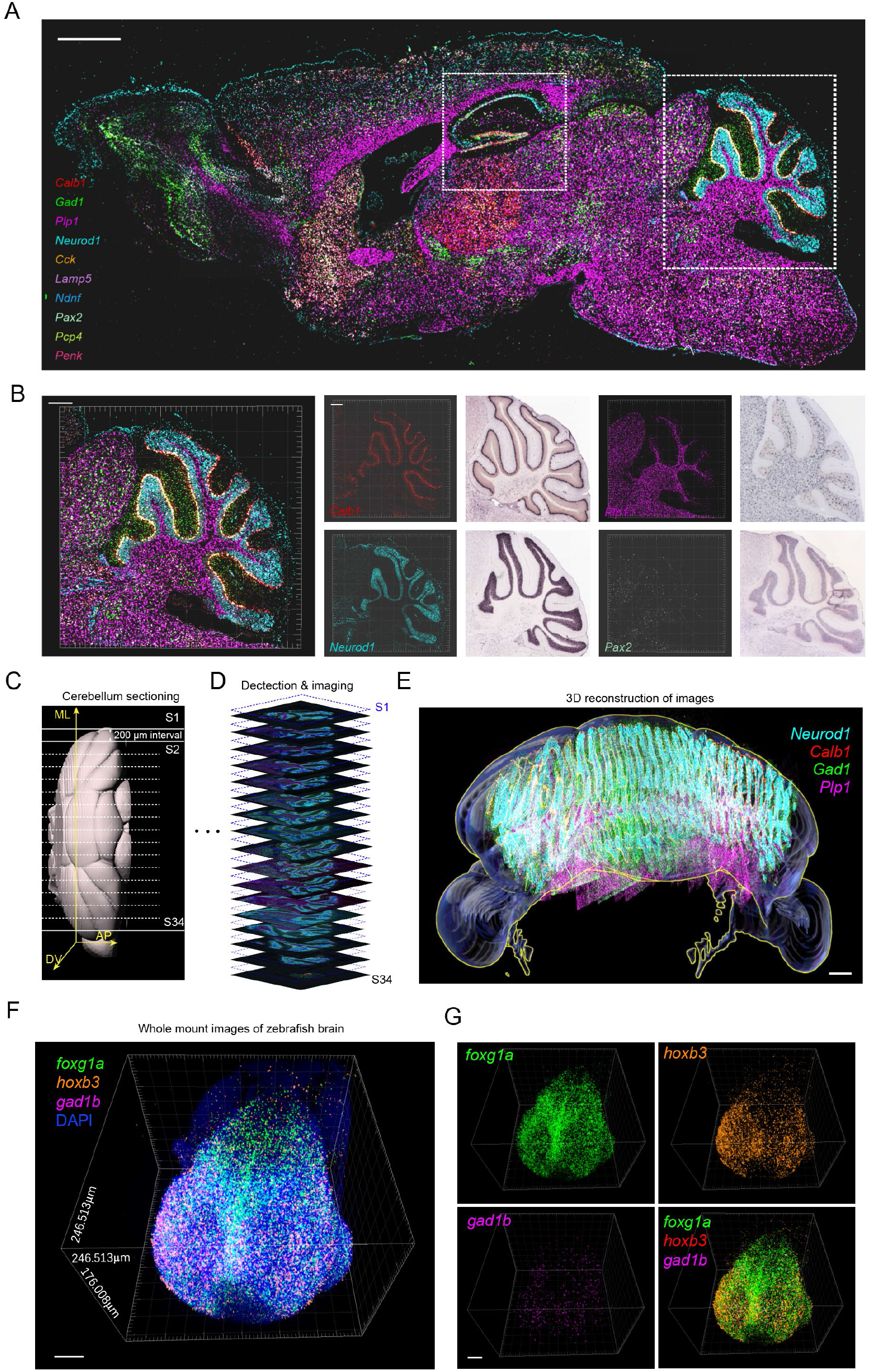
A large-field of view, 3D reconstruction and whole mount spatial profiling of multiple genes. (**A**) Spatial transcriptomic mapping of ten genes (*Calb1, Gad1, Plp1, Neurod1, Cck, Lamp5, Ndnf, Pax2, Pcp4, Penk*) in the whole sagittal section of mice brain. Rectangle region with dashed line in cerebellum and hippocampus were enlarged for view in B and Fig.S5B, respectively. (**B**)Left panels are the enlarged view of rectangle regions in A for ten genes codetection in cerebellum. The gene expression patterns in cerebellum detected by MiP-Seq are consistent with the results from Allen Institute of Brain Science (middle and right panels). (**C**-**E**) Flow chart of 3D reconstruction of fluorescent images for multiplexed genes (*Calb1, Gad1, Neurod1, Plp1*) detection in the whole cerebellum. 34 sagittal sections selected from the whole cerebellum with 200 µm interval were subjected to multiplexed genes detection by MiP-Seq and imaging, followed by reconstruction. (**F-G**) 3D view of the expression and distribution of *foxg1a, hoxb3* and *gad1b* genes detected by MiP-Seq in the whole mount of zebrafish telencephalon. Scale bars, 1000 µm in A, 300 µm in B, 500 µm in E, 30µm in F and G.

**Figure 3.**
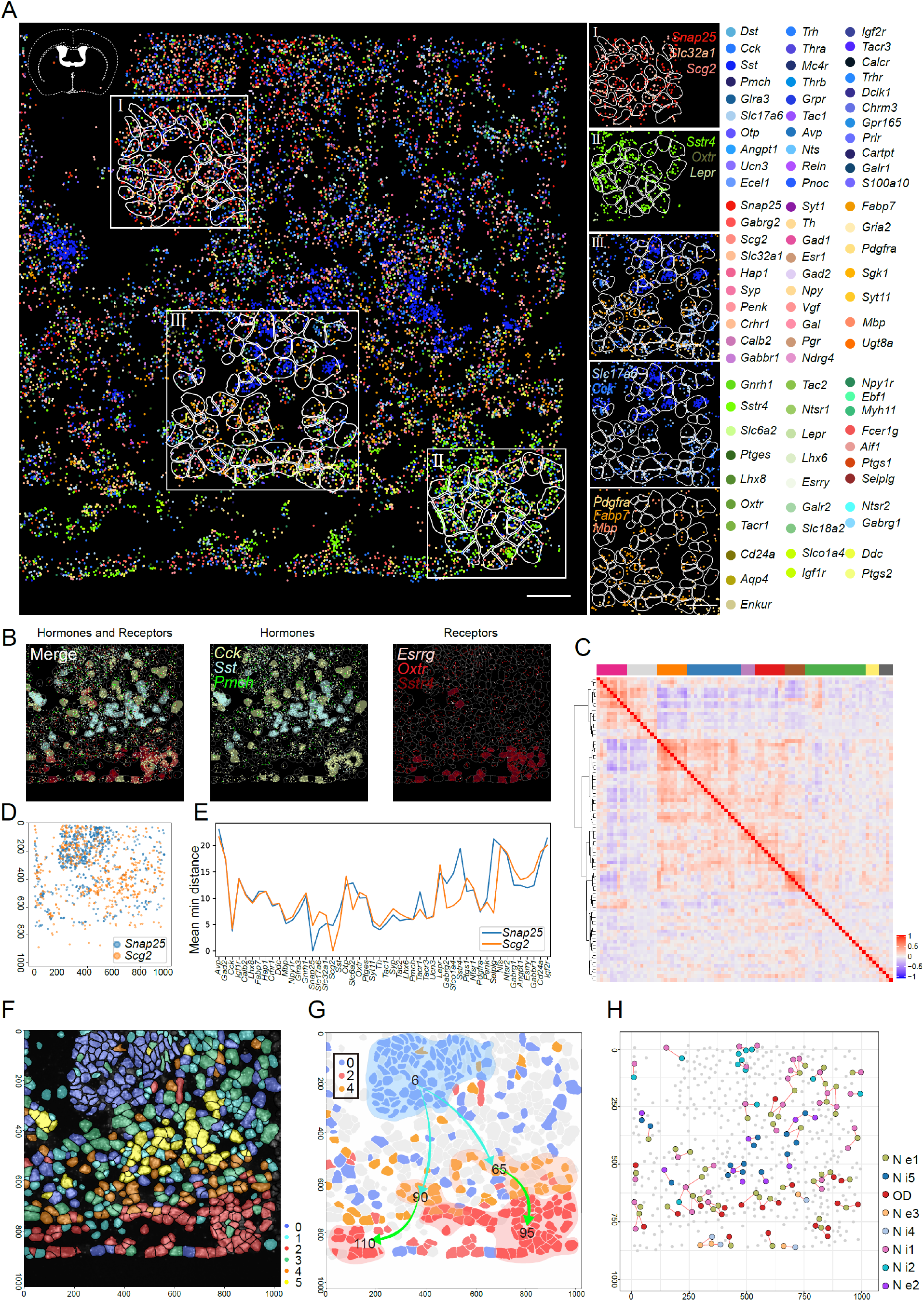
Spatial transcriptomic analysis of 100 genes in nucleus of the horizontal limb of the diagonal band (HDB) by MiP-Seq. (**A**) Spatial transcriptome of 100 genes in HDB detected by MiP-Seq. Transcripts for each gene are pseudo colored. Areas marked with rectangle (I, II, III), which display characteristic spatial-pattern for certain genes transcripts are illustrated separately on the right panel. (**B**) The spatial distribution of hormone-related genes (green) and receptor-related genes (red). When the counts of the corresponding gene in each cell is greater than 10, the whole cell is marked by the color of the corresponding gene. (**C**) Heatmap of top spatial genes co-expression modules with different colors. (**D**) Spatial distance analysis of different genes in HDB. High correlation with the spatial co-expression between gene pairs of *Snap25* and *Scg2*. (**E**) The spatial distance of gene pair (*Snap25* and *Scg2*) to top 50 expressed genes listed in horizontal axis. (**F**) The spatial distribution of six clustered cells in HDB. (**G**) Scatter diagram shows the spatial trajectory inference between cluster 0 and cluster 2 or 4. Two lines and directions indicate distinct differentiation trajectories. The number indicate the subclasses. (**H**) Visualization of *in situ* spatial interaction of ligand and receptor among cells clusters. Lines indicate potential cell pairs that have ligand-receptor interactions. Scale bar, 30 µm in A.

According to the spatial transcriptome profiles, the 100 genes identified in the HDB were clustered into six subclasses (Clusters 0 to 5). And we observed the six subclasses displayed specific spatial distribution pattern along dorsal-ventral axis (Fig.3F and Fig. S8F-S8G). A spatial pseudotime analysis revealed a temporal transition from Cluster 0 to Clusters 2 and 4 (Fig. 3G). As cells of Cluster 0 were obviously clustered together based on their spatial distribution, we characterized the relationship between these cells and other clusters through cell trajectory inference. By combining the spatial information and gene expression profiles, a pseudotime-space analysis revealed that two branches in Cluster 0 (subclass 6) were connected to Cluster 2 (Subclasses 65 and 90) and Cluster 4 (Subclasses 95 and 110). It will be of great interest to investigate the development and correlated functions of each cell cluster in the HDB because these clusters are elegantly organized in different locations.

To further explore the detailed spatial cell-cell interaction network in the HDB, we performed a detailed analysis to classify cells into more subclusters (Fig. S9). Considering the spatial localization of the cell clusters in the HDB, we constructed a spatial cell-cell colocalization network (Fig. S10A-S10D). High-frequency colocalization between oligodendrocytes (OD) and inhibitory neurons in Subclass 4 (Ni4) was observed (Fig. S10D). A spatial cell-cell interaction analysis based on both spatial colocalization and ligand-receptor gene expression information revealed specific ligand-receptor interaction patterns between certain adjacent cell clusters (Fig. 3H). For example, high expression of *Sst* in the inhibitory neuron Subclass 5 (Ni5) and *Sstr4* in the adjacent excitatory neuron Subclass 1 (Ne1), as well as *Tac1* in the inhibitory neuron Subclass 5 (Ni5) and *Tacr3* in the adjacent excitatory neuron Subclass 2 (Ne2) was observed (Fig. S10E). In summary, our *in situ* sequencing results provides a detailed spatial transcriptome in HDB, which may pave the way to understanding the physiology of the HDB at the single-cell spatial level.

Based on high throughput and highly efficient delineation of the spatial profiles of multiplexed genes, MiP-Seq was further utilized to probe dynamic spatial transcriptome changes in response to pathogen infection. To this end, we delineated the spatial transcriptomic alterations of the PVN in response to *Mycobacterium tuberculosis* (*M*.*tb*) infection. The spatial transcriptomic atlas of 217 genes, including cell markers, hormones, receptors, and immune-related genes, was decoded at the single-cell level through three rounds of sequencing (Fig. S11A). The spatial transcriptomic analysis revealed the upregulation of several immune-related genes (*Ly6h, Ccl3*, and *Bcl6b*) and receptor genes (*Gpr165, Calcr*, and *Gria1*) in the PVN upon *M*.*tb* infection (Fig. S11B and data not shown). Meanwhile, the genes *C1qb, Pgm2l1*, and *Ptgs2* were downregulated in the PVN upon *M*.*tb* infection (Fig. S11B). Furthermore, we analyzed differential expression genes (DEGs) in certain cell types based on signal density quantification in the PVN and obtained DEGs in single endothelial cells, astrocytes, microglia, and neurons upon *M*.*tb* infection (Fig. S11C-S11D).

DNA is elegantly folded and organized in the nucleus, and the three-dimensional (3D) genome architecture orchestrates the transcription of genetic code scripts. We therefore applied MiP-Seq to decipher the spatial organization of DNA in brain slices (Fig. 4A). To this end, a padlock probe was first hybridized to a DNA target together with the initial primer after DNA denaturation and ligated using T4 DNA ligase. As shown in Fig. 4B and Fig. S12A, the spatial localization of the *Rbfox3* and *Nr4a1* genes was clearly revealed by MiP-Seq. Furthermore, we utilized MiP-Seq to detect DNA point mutations in C6 glioma (Fig. 4C). Our data unambiguously showed both WT and mutated *Nptxr* genes in glioma cells (Fig. 4C and Fig. S12B).

**Figure 4.**
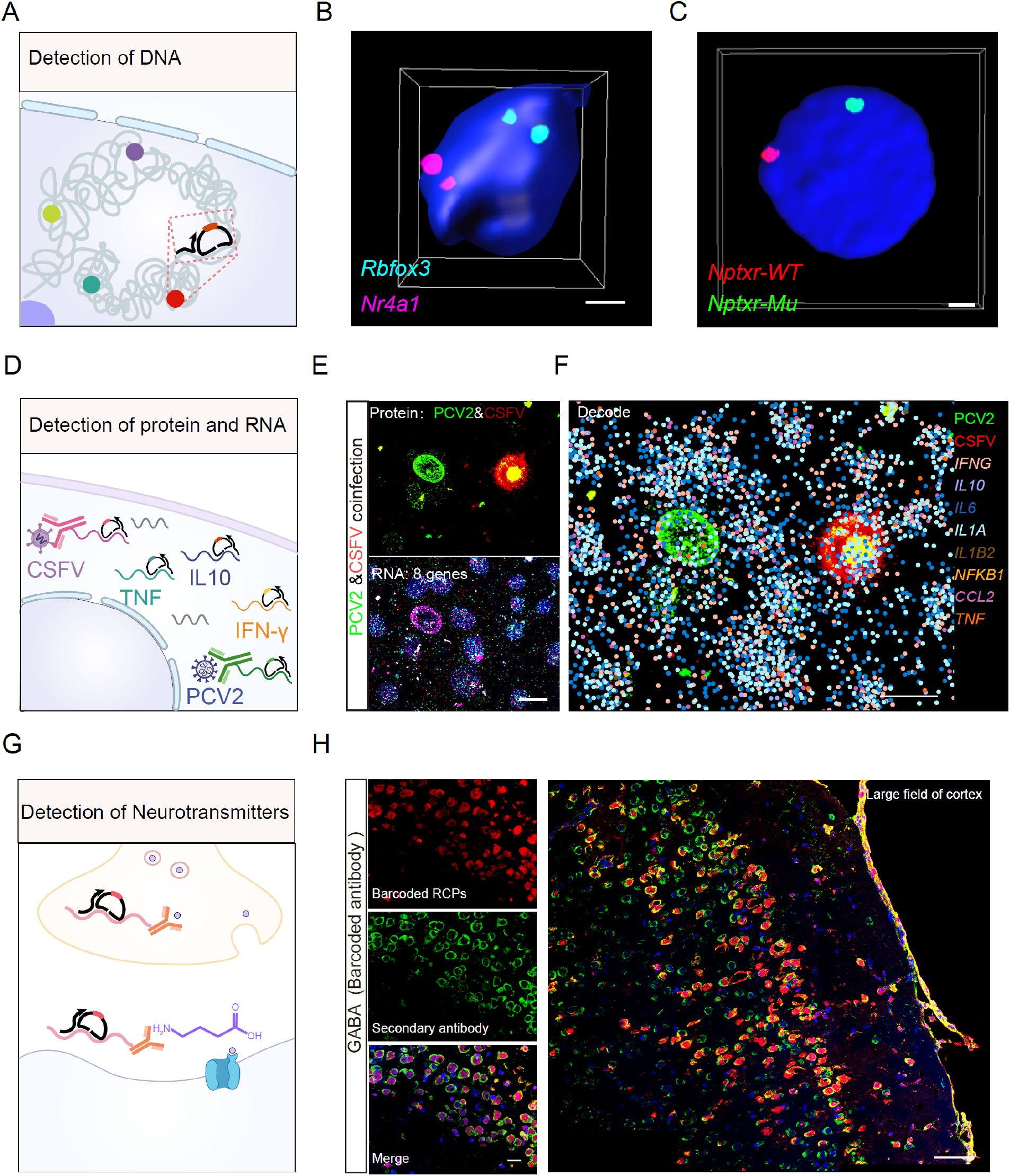
Detection of DNA, RNA, protein, and neurotransmitter by MiP-Seq. (**A**) Schematic diagram of *in situ* co-detection for multiple genomic loci via MiP-Seq. (**B**) The *in situ* detection of *Rbfox3* and *Nr4a1* gene loci located in different chromosomes of nuclei in mouse brain slice by MiP-Seq. (**C**) *In situ* sequencing of *Nptxr* point mutation located in different chromosomes in C6 cell. (**D**) Schematic diagram of simultaneous *in situ* detection of multiple proteins and RNAs. (**E-F**) The mRNAs for eight cytokine or chemokine genes and two virus-specific proteins (E2 for CSFV and Cap for PCV2) in PK-15 cells co-infected by PCV2 and CSFV were simultaneously detected by MiP-Seq. (**G**) Schematic diagram of *in situ* detection of GABA neurotransmitter via MiP-Seq based on nucleic acid-conjugated antibody. (**H**) Identical signals of GABA detected by MiP-Seq with oligonucleotide-conjugated antibodies (Cy3 red fluorescence) to that by classic immunostaining (Alexa Fluor 488 green fluorescence) were observed in the cortex region of mice brain. Scale bars, 2 µm in B and C, 20 µm in E, F and in left image of H, 60 µm in in right image of H.

Furthermore, MiP-Seq was used for *in situ* detection of proteins by *in situ* sequencing of antibody-conjugated nucleic acids. Our data demonstrated the accurate detection of Pol II in PK-15 cells and Orexin in mouse brains by MiP-Seq, and this specificity was validated by traditional immunostaining using the same cells which showed identical gene expression patterns (Fig. S13 and S14A). Notably, MiP-Seq showed much higher sensitivity compared to that achieved with secondary antibody immunostaining due to the strong signal amplification realized through RCA (Fig. S13 and S14A). In addition, we simultaneously detected four proteins, including interferon gamma tagged with Flag, Pol II, the PCV2 cap protein, and the CSFV E2 protein, in PCV2- and CSFV-coinfected PK-15 cells (Fig. S14B), suggesting high-throughput capacity of MiP-Seq for *in situ* protein detection. Furthermore, we performed simultaneous detection of proteins and mRNAs to explore the dynamic expression of 8 cytokine/chemokine genes in response to PCV2 and CSFV virus infection via MiP-Seq (Fig. 4D-4F and S15).

The precise spatiotemporal distribution of biomolecules such as neurotransmitters and hormones play a pivotal role in cell communication and activation. A high-throughput and highly sensitive detection method that leads to subcellular resolution for biomolecules is highly desired^36^. As many biomolecules have semi-antigen characteristic and can be specifically detected by antibodies, we applied MiP-Seq to explore the spatiotemporal distribution of neurotransmitters by *in situ* sequencing of antibody-conjugated nucleic acids (Fig. 4G). To this end, we used γ-aminobutyric acid (GABA), an inhibitory neurotransmitter, for biomolecule detection in brain slices. As shown in Fig. 4H, the spatial distribution of GABA was clearly revealed by MiP-Seq, and this result was validated by traditional immunostaining. Together, these data showed that the versatile MiP-Seq strategy can be employed to detect diverse biomolecules, including DNAs, RNAs, proteins, neurotransmitters, and hormones.

The diverse and specific function of individual cells is highly dependent on their gene profile. In the central nervous system, neighboring neurons can exhibit distinct functional activity and a spectrum of gene expression. By developing an *in vivo-in situ* approach to combine two-photon Ca^2+^ imaging and MiP-Seq in the same animal, we were able to relate the functional activity and gene expression profile in the same individual neurons. Briefly, *in vivo* two-photon Ca^2+^ imaging was first performed in the primary visual cortex of the Thy1-GCAMP6s mouse to identify spike-related somatic calcium signals from all neurons in a volume of approximately 350×350×80 μm^3^. Subsequently, MiP-Seq was performed on the slices in the corresponding imaging region with different neuronal marker genes and neurotransmitter genes (such as *Gad1, Vip*, and *NPY*). The nuclear signals in the *in situ* MiP-Seq images were used to align the *in vivo* two-photon Ca^2+^ image stacks (Fig. 5A-5C and Fig. S16A-S16B). For each identified neuron, the distinct Ca^2+^ activity and gene expression profile were extracted from the Ca^2+^ images and the MiP-Seq dataset, respectively (Fig. 5D and Fig. S16C-S16D).

**Figure 5.**
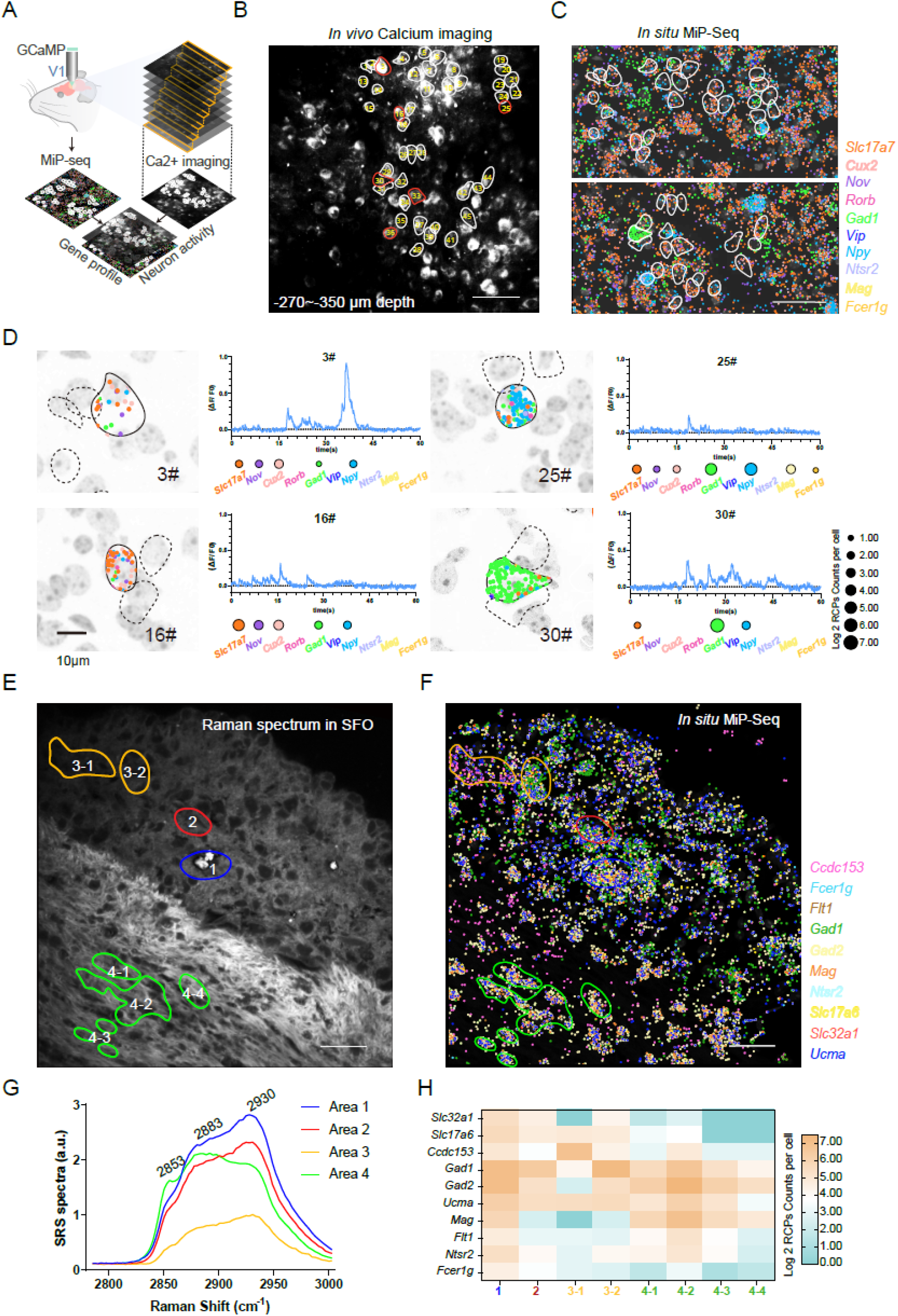
Integrated *in situ* muti-omics of multiple imaging analysis and spatial gene profiles detected by MiP-Seq. (**A**) Schematic diagram for the integrated *in situ* multi-omics of *in vivo* calcium imaging and MiP-Seq to link the transient physiological status and *in situ* gene expression profile in mice visual cortex. (**B**) *in vivo* calcium imaging in visual cortex (−270∼-350 µm depth). Cells marked with circle were numbered (1 to 45) and subjected to *in situ* gene expression profile analysis via MiP-Seq. (**C**) Co-detection of 10 marker genes transcripts in brain slice from the same mice visual cortex corresponding to the calcium imaging area. (**D**) Correlation between gene expression and calcium signal response in cells. (**E**) Hyperspectral stimulated Raman scattering (SRS) of subfornical organ (SFO) on coronal sections (70 µm thickness) of mice brain. Regions with colored circles and numbers indicate distinct anatomical structure. (**F**) Ten cell-marker genes were detected by MiP-Seq in adjacent brain section to E. (**G**) Average SRS spectra of the three chemical species for regions in E. (**H**) Quantification of signals for ten marker genes in regions circled in F by counting RCP number in each region. Scale bars, 50 µm in B-C, 10 µm in D, 30 µm in E-F.

Furthermore, we combined MiP-Seq with hyperspectral stimulated Raman scattering imaging ^37^ to coordinate the biochemical molecule atlas with the *in situ* gene profile. The *in situ* expression of ten cell marker genes was detected in the subfornical organ in mouse brains by MiP-Seq and was aligned to the nearest Raman spectrum images (Fig. 5E-5H and Fig. S17). The specific Raman spectrum and the correlated gene expression pattern are shown in Fig. S17. We also coupled vascular structure images with MiP-Seq data and simultaneously observed the expression of the *Gad1* gene and the microvascular structure in the ARC (Fig. S18).

During multiple rounds of *in situ* high-throughput decoding, optical signal crowding^19^, especially for high-expression level genes, was the most challenging step for signal interrogation. To overcome this problem, we developed a sequential dilution MiP-Seq strategy in which the genes were allocated to different sets of barcodes with different unique dilution primer sequences. In this way, the signals were diluted in individual rounds of imaging by using different sequencing primers. Notably, we selected certain genes as reference genes, which were sequenced in every round by anchor primers (Fig. 6A). After multiround sequencing and imaging, all the gene signals were superimposed based on the alignment of reference gene signals to reconstruct the high-density spatial transcriptomic map. As a proof of concept, two rounds of sequential dilution MiP-Seq for *Cxcl2, Agtr1b, Slc17a6, Ccl2, Il1b, Col1a1, Tnf*, and *Calcr* using reference gene *Gfap* and *Gad2* for alignment was performed. As shown in Fig. 6B-6D, our strategy can indeed successfully realize precise *in situ* gene signal dilution and expression pattern reconstruction. Thus, this sequential dilution MiP-Seq method may greatly facilitate precise interrogation of high-density signals during high-throughput transcriptomic mapping.

**Figure 6.**
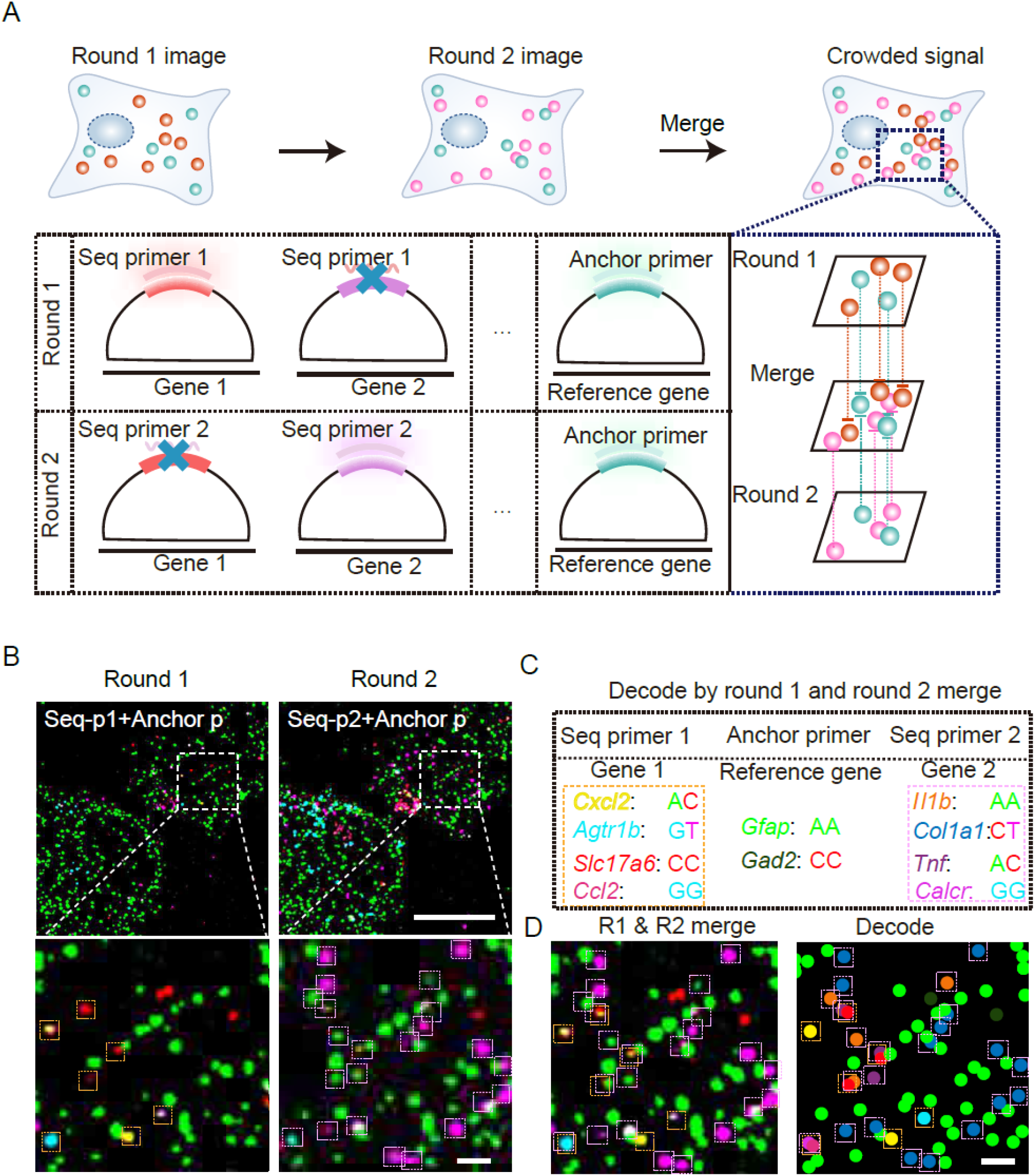
Sequential dilution strategy for optical crowded signals resolving. (**A**) Schematic diagram of sequential dilution strategy for optical crowded signals resolving. Probing highly crowded signals of genes in single cells is diluted into multiple rounds. Fiducial genes are interrogated by anchor primer which are sequenced in each round, while the rest of genes are divided into multiple rounds and sequenced by different sets of sequencing primers. Based on the registration of the fiducial gene signal, the signals of the rest of genes from different sequencing rounds are reconstructed to realize crowded signals interrogation in single cells. (**B-D**) Through two rounds of MiP-Seq in culture neurons from mouse cortex, mRNA signals of *Cxcl2, Agtr1b, Slc17a6* and *Ccl2* (round 1), as well as *Il1b, Col1a1, Tnf and Calcr* (round 2) were reconstructed based on registration of fiducial gene *Gad2* and *Gfap* signals. Scale bar, 20 μm in low magnification of B, 2μm in higher magnification of B and in D.

## Summary and Discussion

In the present study, we developed a high-throughput MiP-Seq strategy to efficiently delineate multiplexed DNAs, RNAs, proteins and neurotransmitters atlases in tissues at subcellular resolution. In contrast to current *in situ* sequencing methods, MiP-Seq leverages a dual-barcode padlock probe and pairwise sequencing strategy, which profoundly increases the decoding capacity compared to that of other methods (10^N^ vs. 4^N^) and requires fewer sequencing rounds. Thus, MiP-Seq can be used to reduce the sequencing and imaging cost dramatically by reducing the sequencing time by approximately 50%. Moreover, the dual-barcode and pairwise sequencing strategy significantly reduces imaging time, which minimizes laser damage, a critical problem during *in situ* sequencing^38-41^. Moreover, our design of an initiator primer and a padlock probe increased the signal specificity and efficacy in comparison to those achieved through current *in situ* sequencing methods. Compared with high-throughput FISH techniques such as MERFISH and Seq-FISH, MiP-Seq can detect short RNAs (with fewer than 500 bp) and can differentiate single base pair mutations, while requiring fewer cycles for signal decoding.

Since MiP-Seq can achieve single-nucleotide resolution, we applied it to detect tumor gene mutations and allele-specific expression of parental genes. As this method can simultaneously detect gene copy number variations and gene mutations in intact pathological sections, it may be extensively used for precision tumor diagnostics. The detection of allele-specific expression of parental genes can greatly improve the understanding of many fundamental biological phenomena, such as heterosis, dominant inheritance, and gene epistasis. Notably, our data showed that MiP-Seq is sensitive to RNA m6A modification and thus exhibits the potential to differentiate m6A-modified RNA from unmethylated RNA. In the future, it will be of great interest to develop a DNA bisulfite switch-like technique to chemically replace the m6A-modified base with another base, which would enable the precise detection of an m6A or other RNA modification *in situ* using MiP-Seq.

Based on the high efficiency of MiP-Seq, hundreds of genes were simultaneously detected in only 2-3 sequencing cycles, which enabled us to map the spatial profile in the HDB and to explore the *M*.*tb* infection-induced dynamic spatial transcriptome alterations in the PVN. These outcomes exemplify the use of MiP-Seq for delineating the spatial omics of certain brain nuclei and identifying alterations in spatial omics at certain locations and in specific cell types. On the basis of the spatial profile of the HDB that was identified by MiP-Seq, many kinds of cell-cell interactions, such as autocrine, paracrine, and long-range interactions between hormones or neurotransmitters, can be identified. It is important to further decipher long-range interactions, especially circuit-based long-range interactions, in the nervous system. In this scenario, the combination of a neural circuit tracing technique and *in situ* detection of ligand-receptor gene expression data will be needed.

Furthermore, in this study, we combined MiP-Seq with *in vivo* Ca^2+^ imaging and Raman imaging to obtain a tissue atlas with multidimensional information, which is highly desired to connect cell phenotypes and functions with gene expression profiles. MiP-Seq can also be combined with other imaging techniques, such as pH imaging, oxidative stress imaging, and electroactivity imaging. Although we did not directly determine the relationships between gene expression pattern with specific calcium dynamics findings at this stage of our research, our method provides a proof of concept to explore the correspondence between a dynamic gene expression profile and distinct cell functions and phenotypes at the *in situ*, single-cell and high-throughput levels. With the advancements in deep learning and the increase in the number of integrated *in situ* genotypes and different kinds of phenotype data, including spatiotemporal biomolecule information and calcium activity, it will pave the way to predict cell function by *in situ* gene profiles, and vice versa, in the future.

Moreover, we successfully performed an *in situ* multiomics analysis based on MiP-Seq to delineate the spatial landscape of DNAs, RNAs, proteins, and small biomolecules (neurotransmitters) at subcellular resolution. This *in situ* multiomics method has not been previously demonstrated. Considering its high-throughput detection capacity, MiP-Seq can be applied to comprehensively decipher the 3D genome landscape by multiple-cycle sequencing, which will greatly increase the understanding of coordinated gene transcription in the 3D genome. For *in situ* detection of proteins and biomolecules, aptamers can be utilized because they can be engineered to specifically bind to proteins/biomolecules and can be easily conjugated to nucleic acids (Fig. S19). In addition, since optical signal crowding causes a “bottleneck” in almost all imaging-based spatial-omics methods, we developed a sequential dilution strategy to solve this problem using a set of sequencing primers and an anchor gene primer. This innovative dilution strategy will greatly facilitate the precise interrogation of high-density signals, such as those of highly expressing genes.

Overall, MiP-Seq is an efficient method for the spatial profiling of 10^N^ genes with only N rounds of sequencing for high-throughput, single-nucleotide, and multiomics detection ability. It can be combined with functional imaging to obtain a tissue atlas with multidimensional information. In this emerging research field, many of these techniques need to be dramatically improved, such as super resolution and low toxicity imaging systems, long-read sequencing strategies, and the integration of functional imaging with *in situ* metabolomic mass spectrometry. The rapid evolution of these techniques will lead to the multidimensional reconstruction of detailed comprehensive molecular and functional maps of many tissues and organs, which may lead us to a new frontier of biomedical research.

## Materials and Methods

### Animals

All mice were kept at 22°C in SPF conditions with food and water available *ad libitum*. Animal care procedures and experiments were approved by the Scientific Ethics Committee of Huazhong Agricultural University, Hubei (HZAUMO-2021-0041, HZAURA-2021-0006), and conducted ethically according to the Guide for the Care and Use of Laboratory Animals of the Research Ethics Committee of Huazhong Agricultural University.

### Establishment of rat C6 glioma model

Wistar rats (170-180 g) were anesthetized by intraperitoneal injection of mixed anesthetic (10% urethane, 2% chloral hydrate, 3mg/mL xylazine) at 0.9 mL/100 g body weight. The head of Wistar rats was fixed on a stereotaxic apparatus. After surgical exposure of the skull, holes were drilled over the sites of the right caudate nucleus of the rat brain. Then, 10 μL of 1.0×10^7^ trypsinized C6 cell suspension in serum-free F12K medium was injected into the right caudate nucleus (AP, 1 mm; ML, 4 mm; DV, 5 mm) with a micro-syringe at speed of 1 μL/min. After injection, the needle was kept for 5 mins to fully deposit the cells. Rat skull was sealed with bone wax and sutured, then rats were returned to the home cage. Two weeks later, the rats were perfused transcardially with 4% PFA in PBS.

### Pups for *in situ* allele specific SNP detection

Female C57BL/6 mice were crossed with male BALB/c mice. The offspring mice were used for *in situ* allele-specific SNP analysis. Partial brain tissues of offspring mice were subjected to transcriptome sequencing. After bioinformatics analysis, multiple allele-specific SNPs were identified for *in situ* allele-specific SNP via MiP-Seq. Brains of offspring mice at P4 were fixed and sectioned for MiP-Seq.

### *Mycobacterium tuberculosis* (*M*.*tb*) infection in mice

C57BL/6 female mice (1.5±0.5 g) at postnatal day 2 (P2) were intraperitoneally injected with either 10^5^ CFU *M*.*tb* H37Ra-BFP or saline control. After 2 hours, pups were transcardially perfused with PBS followed by 4% PFA in PBS. Mice brains were dissected out and post-fixed in 4% PFA for 12 hours at 4°C, then cryoprotected with 30% sucrose in PBS for frozen sections.

### *In vivo* calcium imaging

The male adult C57BL/6J-Tg (Thy1-GCaMP6s) mice were used for *in vivo* two-photon calcium imaging of primary visual cortex neurons activity. Mice were anesthetized with chloral hydrate (0.05 mg/kg) before surgery. A titanium headplate was attached to the skull with dental cement. A craniotomy (4 mm in diameter) was performed above the right primary visual cortex. Dextran Texas red (Thermo Fisher, D3329) was carefully pressure-injected at 3 locations in the craniotomy with a glass pipette, resulting in 3 fluorescent landmarks along the pipette tracks, which were later used to assist alignment of the *in vivo* image stack and post-mortem images for MiP-Seq. Enrofloxacin (1 mg/mL) was injected daily for at least 3 days after surgery. *In vivo* two-photon imaging was performed at least 2 weeks after the surgery.

The activity of cortical neurons was recorded by imaging fluorescence changes with a galvo-resonant scanning microscope (Scientifica) and a mode-locked Ti:sapphire laser (Mai Tai, Spectra-Physics) at 920 nm through a ×16 water immersion objective (0.8 NA, Nikon). Imaging frames of 512 × 512 pixels were acquired at 30 Hz in Scanimage 4.0. To obtain Ca^2+^ activity from all neurons in a cortical volume of approximately 365 × 365 × 80 µm^3^ at the depth of 150-300 μm below the pia, images were recorded at several cortical depths with a spacing of 2.5 µm. After functional imaging, multiple 3D-image stacks covering larger volumes starting from the brain surface were acquired to assist image alignment.

### Cell culture

HEK-293T cells were seeded into 96-well culture plates (PerkinElmer, 6055500) at 9×10^3^ cells/well and cultured in DMEM (Gibco, 11965118) with 10% FBS (Gibco, 10093). For mouse brain microvascular endothelial cell line (BEND3; ATCC CRL-2299) and rat glioma cell line (C6; ATCC CCL-107) culture, 1×10^6^ cells were seeded in a confocal culture dish (BS-15-GJM, Biosharp) coated with 100 µg/mL poly-D-lysine (Sigma, P0899) and maintained in DMEM with 10% FBS. 1×10^6^ human brain microvascular endothelial cell (HBMEC, ATCC CRL-3245) were seeded in a confocal culture dish with DMEM containing 10% FBS, 2 mM L-glutamine (Gibco, 25030081), 1 mM sodium pyruvate (Gibco, 11360070), 1× MEM nonessential amino acids (Gibco, 11140050), 1×MEM amino acid solution (Gibco, 11130051), 1×MEM vitamins (Gibco, 11120052), and 1×penicillin-streptomycin. The neocortex of mouse embryos at E15-16 was dissected out and digested by 12.5 mg/mL trypsin (Gibco, 27250018) into individual cells. The cells were washed twice in DMEM containing 0.001 U/µL DNase I (Takara, 2270A) and 5% FBS, then washed once by DMEM containing 5% FBS. Next, the cells were seeded in a confocal culture dish coated with 100 µg/mL poly-D-lysine (Sigma, P0899) at a density of 2×10^5^ cells per dish. After 8 hours, the medium was replaced by Neurobasal (Gibco, 21103049) which contains 1.25% FBS, 2% B-27 (Gibco, 17504044) supplement, 0.5 mM L-glutamine (Gibco, 25030081), and 0.5% Penicillin-streptomycin (Gibco, 15140163), and cultured for 7 days. All above cell lines were incubated at 37 °C, 5% CO_2_.

For *in situ* m6A methylation detection via MiP-Seq, commercially synthesized RNA with m6A methylation (Sequence: GGAAAGGA(m6A)CUAAAAA) and the corresponding unmethylated RNA were transfected into HEK-293T cells by lipofectamine 2000 (Invitrogen, 11668019), respectively. Three hours after transfection, cells were subjected to MiP-Seq.

To prepare cell samples, cells were quickly washed twice with PBS and fixed with 4% paraformaldehyde (BOSTER, AR1069) for 10 mins at room temperature. After fixation, cells were washed three times with PBSTR (PBS containing 0.1% Tween-20 (Promega, 0000395426), 0.1 U/µL RRI (Vazyme, R301-01)) at 5 min each time. The cells were then added with methanol (Sigma, 34860) pre-cooled at -20°C and kept at -80°C for 15 mins. Finally, the cells were sealed by Secure-Seal hybridization chambers (Grace, 621505) for the following MiP-Seq.

### Brain slices and zebrafish brain preparation and pretreatment

Mice and rat brains were dissected out after perfusion with 4% PFA in PBS and post-fixed in 4% PFA for 12 hours at 4°C, then cryoprotected with 30% sucrose in PBS. After embedding in OCT at -80°C, brain sections (8 µm or 10 µm) were cut with a Leica cryostat. All brain slices were stored at -80°C. Before MiP-Seq, brain slices were sealed by Secure-Seal hybridization chambers and pretreated by fixation and permeabilization. Briefly, brain slices sealed by Secure-Seal hybridization chambers were washed twice with pre-cooled DEPC-PBS and fixed with 4% PFA for 10 mins, then washed three times with DEPC-PBSTR (PBS with 0.1% Tween-20, 0.1 U/µL RRI) for 5 mins each time. Next, slices were added with pre-cooled methanol and kept at -80°C for 15min. Further, slices were treated with 2 mg/ml pepsin (Sigma, P0525000) in 0.1 M HCl at 37 °C for 90 s. After three times washing with DEPC-PBSTR, the brain slices were ready for MiP-Seq.

Zebrafish brains were fixed with 4% PFA in PBS for 12 hours at 4°C and cryoprotected with 30% sucrose in PBS, then followed by two rinses with pre-cooled DEPC-PBS and dehydration with a series of methanol (25% (vol/vol), 50% (vol/vol), 75% (vol/vol) methanol in PBS, and 100% methanol, 5 min in each). After that, zebrafish brain can be stored with 100% methanol at -20°C for several months. Before MiP-Seq, zebrafish brains were rehydrated by a series of methanol (75% (vol/vol), 50% (vol/vol), 25% (vol/vol) methanol in PBS, 5 min in each) and washed three times with DEPC-PBSTR. Then, zebrafish brains were permeabilized with proteinase K (20 mg/ml) (NEB, P8107S) at room temperature for 30min, followed by three times washing with DEPC-PBSTR. Next, the brain was ready for MiP-Seq.

### MiP-Seq probe design

Probes for MiP-Seq included the pair of padlock probes with RCA initiator primer probe for target-dependent RCA, detection probes for RCA, and *in situ* sequencing probes. The probes were designed as follows: (1) Use the self-developed probe searching program to screen the target sequence; (2) Try to choose the exon close to the 5’ end, and select 2 target sites for each gene; (3) Both ends of the padlock probe are complementary to the target sequence (12-16 bp), and the middle sequence is composed of a complementary sequence for anchor primer and a barcode sequence; (4) The 5’ end of the RCA initiator primer is complementary to the 3’ end of the targeting sequence (12-16 bp), and the 3’ end is complementary to the padlock probe; (5) The detection probe (cy3-p4) were used to test RCA efficiency; (6) The probes for *in situ* sequencing have consisted of three anchor primers (anchor-1, anchor-2, and anchor-3) and four query probes (seq-1, seq-2, seq-3, and seq-4) conjugated respectively with Alexa Fluor 488, Alexa Fluor 546, Alexa Fluor 594, and Alexa Fluor 647. All probe sequences are listed in Table S1.

### MiP-Seq for *in situ* RNA detection

Padlock probes were phosphorylated at 5’ end with T4 Polynucleotide Kinase (Vazyme, N102-01) before MiP-Seq. Our MiP-Seq mainly included six sequential steps: annealing of probes to targets, ligation of padlock probes, rolling circle amplification, multiple rounds of sequencing by ligation, imaging, and decoding. In detail, phosphorylated padlock probes (30 nM/probe) and equimolar RCA initiator (30 nM/probe) in hybridization buffer (2X SSC (Sangon, B548109) with 10% formamide (Sangon, A100606) and 20mM RVC (Beyotime, R0108)) were added to the hybridization chamber of cultured cell samples or pretreated brain slices and incubated overnight at 37°C. After annealing of probes to RNA targets, samples were washed twice with DEPC-PBSTR (0.1% Tween-20, 0.1 U/µL RRI in DEPC-PBS) for 20 mins each time, followed by washing with 4X SSC in DEPC-PBSTR once. The ligase reaction mixture (1 U/μL SplintR ligase (NEB, M0375L), 1X buffer, 0.2U/μL RRI) was then added to the hybridization chamber for 2 hours incubation at 37°C. Following ligation, samples were washed with DEPC-PBSTR twice for 20 mins each time and incubated with rolling circle amplification reaction solution (1 U/μL Phi29 (Vazyme, N106-01), 1X RCA buffer, 0.25 μM dNTP, 0.2 μg/μL BSA, 5% Glyceryl) at 30°C for 2 hours. After double washing with DEPC-PBSTR (20 min each time), the samples were ready for sequence by ligation or direct detection by the fluorescent probe.

Next, sequencing mixture (0.2 U/μL T4 DNA ligase (ThermoFisher Scientific, EL0012), 1X T4 DNA ligase buffer, 1X BSA, 1 μM anchor primer, 1 μM query primer) was added to the chamber for 2 h incubation at 25°C. The samples were then washed three times with 10% formamide in 2X SSC for 5 mins each time, and counter-stained by DAPI. Samples were immersed in imaging buffer (10% (w/v) glucose, 40 μg/mL catalase (Sigma C30), 0.5 mg/ml glucose oxidase (Sigma G2133), 0.02 U/µl Murine RNase inhibitor, and 50 mM Tris-HCl pH 8 in 2X SSC) and imaged by Leica TCS SP8 confocal microscope with 60× water lens (NA 1.2). Channels of DAPI, Alexa Fluor 488, Alexa Fluor 546, Alexa Fluor 594, and Alexa Fluor 647 were scanned. After each round of imaging, signals were stripped by stripping buffer (60% formamide in 2X SSC) twice at room temperature for 10 mins each. The sequencing mixture for the next round of sequencing was then added to perform multiple round sequencing for high throughput readout. Images were analyzed via self-developed multi-channel identification, 3D, multi-round proofreading process. Each channel signal was filtered, identified, and registered. Each round of barcodes was registered, and the acquired signals were quality-controlled and tested.

For single-gene quantitative detection, fluorescent probe CY3-P4 (100 nM in 2X SSC), which is complementary to RCP, was added to the hybridization chamber and incubated for 4 hours at 37°C. After three times washing with PBST (PBS with 0.1% Tween-20) for 5 mins each, cell samples were imaged with Leica TCS SP8 confocal microscope with 60× water lens. Images were analyzed by a self-developed pipeline of cell segmentation and counting process for signal dots decoding, counting, and statistical distribution analysis.

### MiP-Seq for *in situ* DNA detection

Cell samples were rinsed twice with PBS and dehydrated in a series of 70%, 80%, 90%, and 100% ethanol for 5 min each. The cells were washed in PBST twice for 2 min each time, followed by incubation in 0.1M HCl for 5 min and two washes in PBST. To digest RNA, cell samples were treated with 50 µl solution consisting of 2X SSC, 100 µg/ml RNase A (Thermo Fisher, EN0531) for 2 h at 37 °C, and then washed three times in 2X SSC at room temperature (2 mins each time).

To denature DNA, cell samples were incubated with 70% formamide in 2X SSC at 85°C for 10min, and then quickly transferred to a series of 70%, 80%, 90%, and 100% ethanol for 5 mins each. Samples were air-dried for 5 mins. After that, 100 nM padlock probe and 100 nM initiator primer in hybridization solution (10% formamide in 2X SSC) were added to the samples for overnight incubation at 37°C. After hybridization, samples were washed once with 25% formamide in 2X SSC and three times with PBST (2 mins each time) at room temperature. The padlock probes on the DNA target were then ligated by 0.2 U/μL T4 DNA ligase (Thermo Fisher, EL0011) in 1X ligation buffer with 0.2 μg/μL BSA for 3 h at 16°C to form an RCA template. After ligation, samples were rinsed in PBST three times at room temperature (2 mins each time) and followed by RCA (1 U/μL Phi29, 1X Phi29 buffer, 0.25 μM dNTP, 0.2 μg/μL BSA, 5% Glyceryl and 2 μM dUTP) at 30°C for 6 h. After washing in PBST twice, cells were incubated with 100 nM of each corresponding detection probe in 2X SSC with 10% formamide at 37 °C for 1 h, then immersed in the anti-quenching buffer for imaging.

### Conjugation of nucleic acid to Antibody

Antibodies of GABA (Invitrogen, PA5-32241), Orexin (Cell Signaling Technology, 16743S), Pol II (ABcam, ab5095), Flag (Proteintech, 66008-3-Ig), CSFV E2 (generous gift by Wuhan Keqian Biological), and PCV2 Cap (generous gift by Wuhan Keqian Biological) were dialyzed in PBS with a 10 kd Slide-A-Lyzer mini dialysis tube (Thermo, 69570) overnight at 4°C, and concentrated to 50-100 μL by Amicon spin filters (Merck, MRCF0R030) at a concentration higher than 1mg/ml. Then, ten times molar amount of NHS-PEG4-Azide (Thermo, 26130) was added to the antibody solution and shaken for 2 hours at 4°C, followed by dialysis in PBS with stirring on ice for 4 hours. After dialysis, the antibody was added with four times molar amount of DBCO-modified oligonucleotide and incubated with shaking at 4°C for 12 hours. Finally, Protein A/G magnetic beads were used (Yepsen, 36417ES03) to purify the antibody conjugated with oligonucleotide.

### Co-detection of protein and RNA by MiP-Seq

1× 10^4^ cells/ml of PK-15 cells (generous gift by Wuhan Keqian Biological) seeded in a confocal culture dish were simultaneously infected with PCV2 and CSFV at 10^2^ TCID50. Cells were treated with D-glucosamine at 37°C for 1 h, then cultured for 24 hours. Subsequently, the supernatant was discarded and cells were fixed with 4% PFA in PBS for 10 mins at room temperature, followed by the procedure of MiP-Seq.

### Co-detection of four proteins during virus co-infection

A confocal culture dish was inoculated with 1×10^4^ cells/ml PK-15 cells, and the AAV-IFNG-flag plasmid was transfected into cells 8 hours after seeding, following the protocol of the lipofectamine 2000 transfection kit (Invitrogen, 11668019). 8 hours after transfection, the cells were infected with 10^2^ TCID50 of PCV2 and CSFV for 12-24 hours. The cells were treated with 300 mM D-glucosamine at 37°C for 1 hour and cultured for 24 hours. Then cells were fixed with 4% PFA in PBS at room temperature for 10 mins for MiP-Seq.

### Hyperspectral stimulated Raman scattering (SRS) imaging

The brain slices from 8-week-old C57BL/6 male mice with 50 μm thickness were imaged by hyperspectral SRS microscopy. Both pump and Stokes beams were set at 30 mW, and the pixel set was 400 × 400 pixels with a dwell time of 10 μs/pixel. For high-resolution imaging, a water immersion objective (N.A. 1.2, UPLSAPO 60× W) was used. The pump and Stokes lasers were set at 800 nm (40 mW) and 1040 nm (100 mW), respectively. The dwell time for NIR SRS imaging was 10 μs/pixel, with a field of view of 250 × 250 μm^2^ with 1600 × 1600 pixels.

### Vasculature structures imaging

After MiP-Seq, the brain sections were incubated with 25 µg/ml DyLight 488 (λ Ex 493 nm; λ Em 518 nm) Lycopersicon Esculentum Lectin (Vector Laboratories) at room temperature for 30 mins for blood vessels staining. Images were acquired using Leica TCS SP8 confocal microscopy with a 405 diode, Alexa Fluor 488, Alexa Fluor 594, and white light laser by 60× water-immersed objective (NA 1.2).

### Image registration

The pipeline for image registration was divided into three sequential steps. Firstly, images along Z-axis (*Z resize*) were resized. The shape of the original image is (1024,1024,*z*_*i*_), where *z*_*i*_ is the z-axis length of the image of round i. To align images of different sequencing rounds, all images from each round were interpolated to the same z-axis length: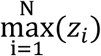. The <u>skimage.transform.resize</u> function from the scikit-image^42^ package was used for interpolation. The image of the first round was selected as the reference cycle. The images from the second and third rounds were aligned and transformed to the first cycle image. Before registration, the <u>white_tophat</u> function (from scikit-image, with parameters: structure_element=“ball”, size=1) was used to improve the signal-to-noise of the images. The registration was performed by two steps: (1) using z-axis projected and gaussian blurred (sigma=5) images as inputs to eliminate the bias in a horizontal direction. (2) using the whole 3D images as inputs to eliminate the bias in the 3D space. The SimpleElastix toolkit ^43^ was used for registration. The ‘affine’ parameter was used during the whole process.

### Candidate spots calling

Firstly, the <u>white_tophat</u> (parameter same to above) was used to improve the signal-to-noise of the registered images. To balance the intensity of all channels, reference spots were identified using the <u>h_maxima</u> function (from scikit-image). The h parameter was set to the 95% quantile intensity of the corresponding channel. Then each channel with the mean image intensity at these reference spots was scaled. The mean image of these normalized channel images was used for further analysis. The <u>h_maxima</u> function was used for candidate spot calling. The h parameter was set to the 95% quantile intensity of the mean image.

### Decoding

The single channels were combined to analyze the intensity of the dual barcode spots. The intensity of the combined channel (ch1 and ch2) is: *l*_*ch*1,*ch*2_ = *sqrt*(*l*_*ch*1_ ∗ *l*_*ch*2_). For each spot, a 5×5×3 (x, y, z) cube was cropped around it. Then the cube was projected to get a 5×5 square matrix. The max intensity point *m*_*i*_ was identified in this matrix. The mean intensity of a 3×3 sub-matrix around the *m*_*i*_ in all 3 rounds and 4 channels were calculated as the representative value *p*_*ij*_(*i* ≤ 3, *j* ≤ 4).

The candidate spots need to be filtered out when: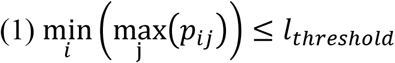, where *l*_*threshold*_ is the threshold value of the intensity; (2) The spot is located on the border of the image. For each round of images, let *VS* = {*s*_1_, *s*_2_, *s*_3_, *s*_4_} as the sorted representative value. In Fig.2, the spot will be considered as the single-channel signal if *s*_1_ > 2 ∗ *s*_2_, otherwise be treated as dual-channel merged signal. In Fig.S11, the signal-to-noise ratio for single-channel 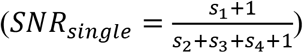 and dual-channel 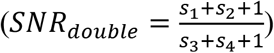 spots were calculated and compared. If SNR _*single*_ > SNR _*double*_ the spot will be recognized as single-channel signal, otherwise as dual-channel signal. For each spot, all the codes from each round were combined to generate their barcode. The gene can be deciphered by matching the barcode with the gene codebook.

All detailed efficiency comparison data are listed in Table S2. And all processed intermediate data are listed in Table S3.

### Cell segmentation and genes assignment

Firstly, the images were projected along Z-axis to get 2D images. To highlight the boundary of the cells, exposure adjustment of the images was performed using the <u>adjust_sigmoid</u> function (from the scikit-image package, with parameters: cutoff=0.03, gain=100), followed by cell segmentation using the cellpose software^44^. For the cells with segmentation difficulty by cellposs, the labelme software was used to perform cell segmentation manually. When the above steps are completed, all cell mask and position information are available. To assign gene transcripts to cells, we calculated the Euclidean distance between *g*_*i*_ and its nearest cell *C*_*i*_ = *argmin*_*j*_ (*D*_*ij*_) for each gene *g*_*i*_. If *D*_*ij*_ ≤ *D*_*threhold*_and assigned *g*_*i*_ to *C*_*i*_, the *g*_*i*_ ‘s position was treated as a part of the pixel of *C*_*i*_. This process was repeated until there was no new assignment occurred.

### Image processing pipeline for low throughput MiP-Seq (when genes ≤ 10)

Another pipeline was used for image processing when the gene number was less or equal to 10. The image registration is not needed as there is only one imaging round. The candidate spots were called on each channel’s image. Before spot calling, white_tophat (structure_element=“ball”, size=1) function was applied for improving the signal-to-noise of the balanced images. Then the <u>h_maxima</u> was used for spot calling. After that, all spots were decoded according to Euclidean distance to its nearest neighbors’ spots and their channel information. A spot was considered as the dual barcode spot when it contained two fluorescence signals within a certain radius. The corresponding genes of the spots were identified according to their channel information and gene codebook. Finally, cell segmentation and gene to cell assignment were performed using the same method as the high-throughput pipeline.

### Cell clustering and co-localization analysis

For cell clustering of HDB nuclei, the count matrix was calculated from the gene to cell relationship after gene to cell assignment. The empty cell contains no genes are filtered out. Then Leiden clustering was performed on the expression matrix, and parameter resolution=1 and resolution=0.6 were used to produce 11 and 6 cell clusters, respectively. The Leiden function of the scanpy ^45^ package was used here. The clustering result was visualized with the uniform manifold approximation and projection (UMAP) embedding. UMAP-learn python package was used here ^45^.

For cell clustering of PVN, we first screened microglia and oligodendrocytes by marker genes before unbiased clustering as the number of them were too small to be clustered. Subsequently, stLearn was used to perform QC (min_genes=5, min_cells=2), integration and unbiased clustering (resolution=0.7) based on the expression matrices of the remaining cells, resulting in total 7 cell clusters (including microglia and oligodendrocytes). The marker genes of the 7 cell clusters were displayed by a heatmap. Merging clusters of the same type led to 5 major subpopulations ultimately, including astrocytes, neurons, AVP-neurons, endothelial cells, and ependymal cells. We then extracted the processed expression matrices from stLearn and performed between-group differential analysis using Seurat (cut off pval≤ 0.05 & |logFC| ≥ 0.3).

For gene co-localization analysis, the image was split into 20×20 chunks and the gene occurrence were calculated in these chunks. Then the correlation between all gene pairs was calculated using the <u>scipy.stats.kendalltau</u> function. To measure the proximity between different cell-types, the adjacent network was constructed (FigS9A). In this network, one node represents a cell, and the edge between two nodes means these two cells were adjacent in space. Then the adjacent matrix of all cell-types was constructed by counting the edge numbers of each node pair type.

### Spatial gene co-expression, cell type interaction, ligand-receptor cell-cell communication, and Pseudotimespace analysis

Giotto v1.0.4 ^46^ was used for the spatial gene co-expression, cell-type interactions, and ligand-receptor cell-cell communication analysis. In detail, the raw matrices were normalized and adjusted with default parameters, and the previous classification results (scanpy, resolution=1) were imported. Next, a “Delaunay network” was created to calculate the cell neighborhood distribution. On this basis, spatial gene co-expression patterns were then identified using “binSpect” function. The analysis of cell-type interactions was performed with “cellProximityEnrichment” function. After filtering the ligand-receptor gene-pairs with Giotto’s database, “spatCellCellcom” function was used to calculate the significant ligand-receptor interactions (cut off p.adj ≤ 0.05 & |log2fc|> 0.1 & lig_nr ≥ 2 & rec_nr ≥ 2).

StLearn v0.3.1 ^47^ was applied to the analysis of spatial trajectory reference. Based on the spatial information and previous broad clusters (scanpy, resolution=0.6), stLearn found 131 sub-clusters (0-130) which were spread across two or more spatially separated locations. The right bottom corner cells of cluster 0 were selected as the root. Then, “st.spatial.trajectory.pseudotimespace_global” function was used to reconstruct the global level spatial trajectory. The figures were generated by Giotto and stlearn or redrew using R packages igraph v1.2.6 ^48^ and ggplot2 v3.3.5 ^49^.

## Acknowledgments

We would like to thank Prof. Fang Yang (Huazhong Agricultural University) for helping in tissue processing; Zhe Hu, Yunguang Li, and Chunmei Shi in Core Facility at HZAU for technical support in imaging; Zhihui Zhang (Spatial FISH Co., Ltd) for technical support in FISH; Xuehan Li and Jiaming Li for helping in RCA; Jie He, Hui Zhang and Mengmeng Jin (State Key Laboratory of Neuroscience, Institute of Neuroscience, Center for Excellence in Brain Science and Intelligence Technology, Chinese Academy of Sciences) for providing zebrafish brain and related help; Wuhan Keqian Biology Company for providing PCV2 and CSFV virus.

## Author contributions

G. C., J. D., and X. W. conceived and designed the project. X. W. performed probe library design and preparation, cell and tissue experiment processing and sample preparation, *in situ* sequencing experiments, imaging, data analysis. W. X., C. W., and Y. Z. performed probe design software development, spatial gene decoding pipeline design, and analysis. L. D. performed the spatial trajectory and spatial Delaunay network analysis. Y. L. performed C6 glioma model establishment, Raman imaging analysis, vascular imaging experiment. Z. W. performed gene *in situ* verification in brain tissue, parental orthogonal experiment, methylation *in situ* detection, *in situ* hybridization of Raman tissue slice experiment. L. S. and G. D. performed conjugation of nucleic acid to antibody and *in situ* detection of proteins. A. G. performed MiP-Seq and HCR3.0 efficiency comparison experiments. H. W. performed MiP-seq *in situ* DNA detection. X. Y. performed *in vivo* calcium imaging and analysis. K.Y. performed RNAseq library sequencing for C6 cells and offspring brain tissue of parental orthogonal. L. Z. and L. Y. performed neuron culture and efficiency verification. Z. L. and D. L. provided bioinformatical analysis. M. L. performed early experimental exploration. G. C., J. D., Y. H., Z. F., P. W., X. W., W. X., and L. D. wrote the manuscript. All authors read and approved the final manuscript.

## Data availability statement

Raw and processed RNA-Seq data have been deposited in GEO (GSE190219). The source image data for this study are available at Zenodo (https://zenodo.org/record/6299029#.Yho8Zd9BxjU). Publicly available datasets used in the study includes GSE113576 and GSE74672.

## Code availability statement

The custom written package and scripts used in this study are available at https://github.com/GangCaoLab/InnerEye.

## Supplementary Figures for

**Fig. S1.**
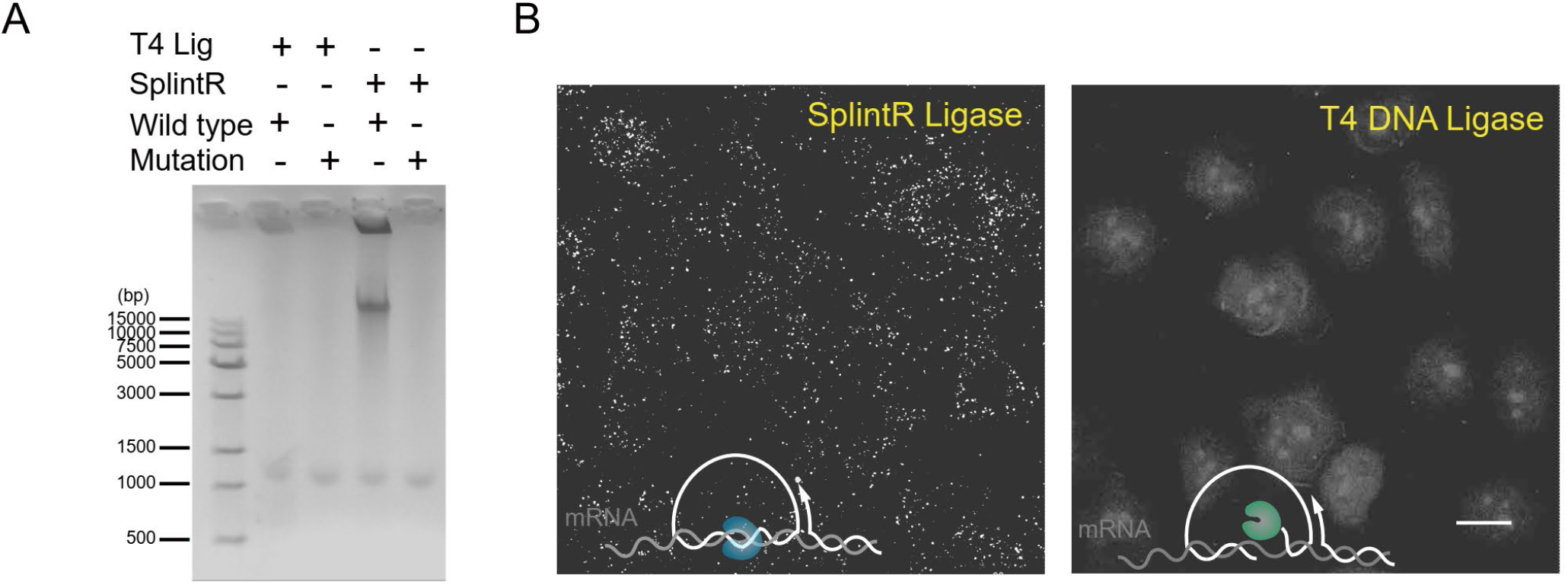
High fidelity and efficiency of RNA-dependent DNA ligase SplintR. (**A**) Gel electrophoresis result for the detection of human wild-type *ACTB* transcripts with wild-type probe (the sequence is fully complementary to the transcript) and mutant probes (single unmatched base adjacent to the ligation point) by T4 and SplintR ligase respectively. (**B**) *In situ* detection of *ACTB* mRNA in HBMEC cells using SplintR ligase (left) and T4 ligase (right). Scale bar, 15 μm in B.

**Fig. S2.**
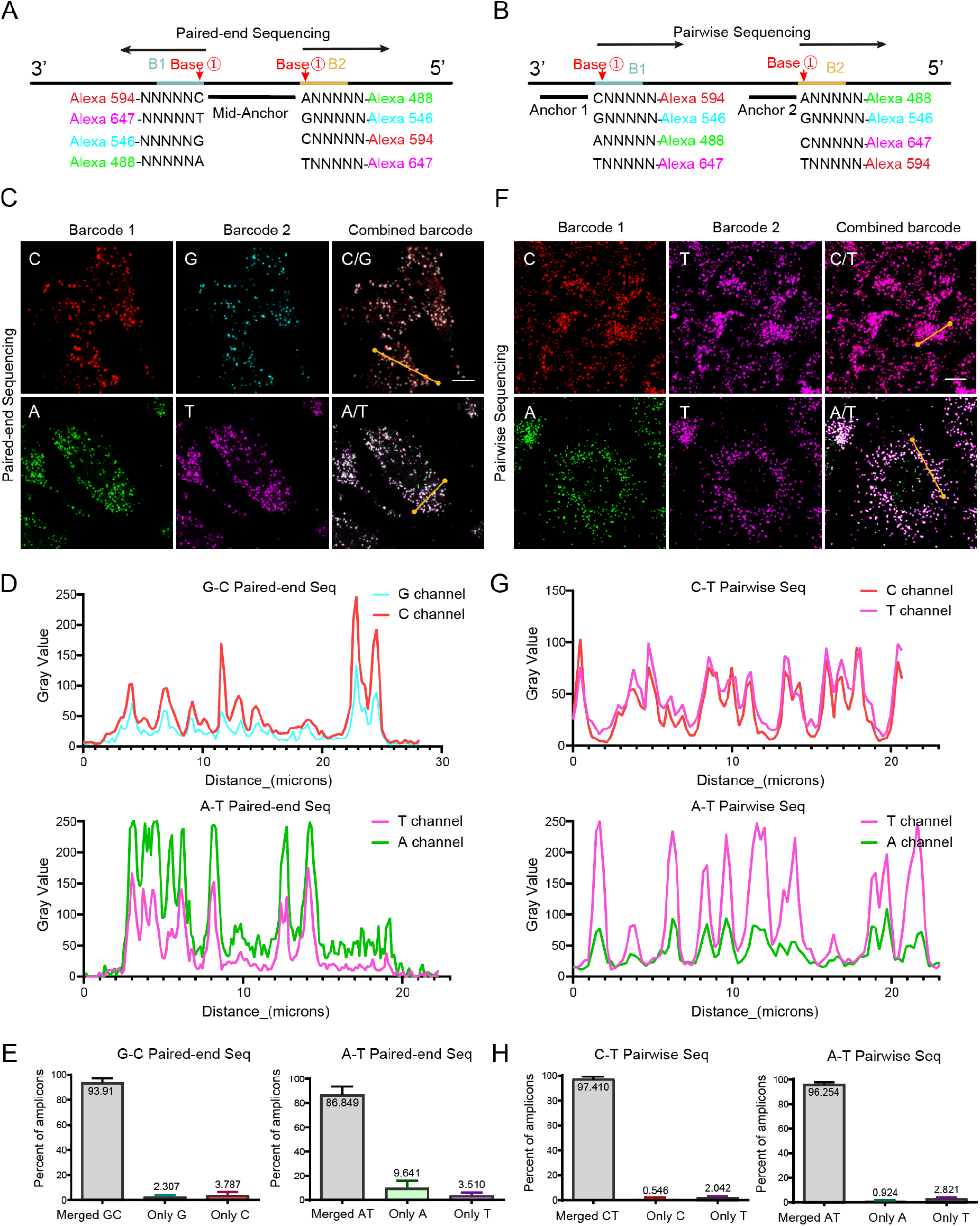
Signal merging of paired bases during dual-barcode sequencing. (**A**) Schematic diagram for paired-end sequencing. The upper and lower sequencing primers are simultaneously ligated to the same mid-anchor primer to interpret double bases of dual barcodes. (**B**) Schematic diagram for pairwise sequencing, in which two sequencing primers are individually ligated to two separated anchors to realize double bases interrogation. (**C**) *In situ* detection of *UBC* (G/C dual bases) and *GAPDH* (A/T dual bases) gene transcripts in HBMEC cells *via* paired-end sequencing. (**D**) Consistence of fluorescent intensity curves for paired G/C bases and paired A/T bases along the yellow straight line in image C, respectively. (**E**) The percentage of merged signals for dual bases during paired-end sequencing. (**F**) *In situ* detection of *UBC* (C/T dual bases) and *GAPDH* (A/T dual bases) gene transcripts in HBMEC cells via pairwise sequencing. (**G**) Consistence of fluorescent intensity curves for paired C/T bases and paired A/T bases along the yellow straight line in image F, respectively. (**H**) The percentage of merged signals for dual bases during pairwise sequencing. Scale bars, 10 µm in C and F.

**Fig. S3.**
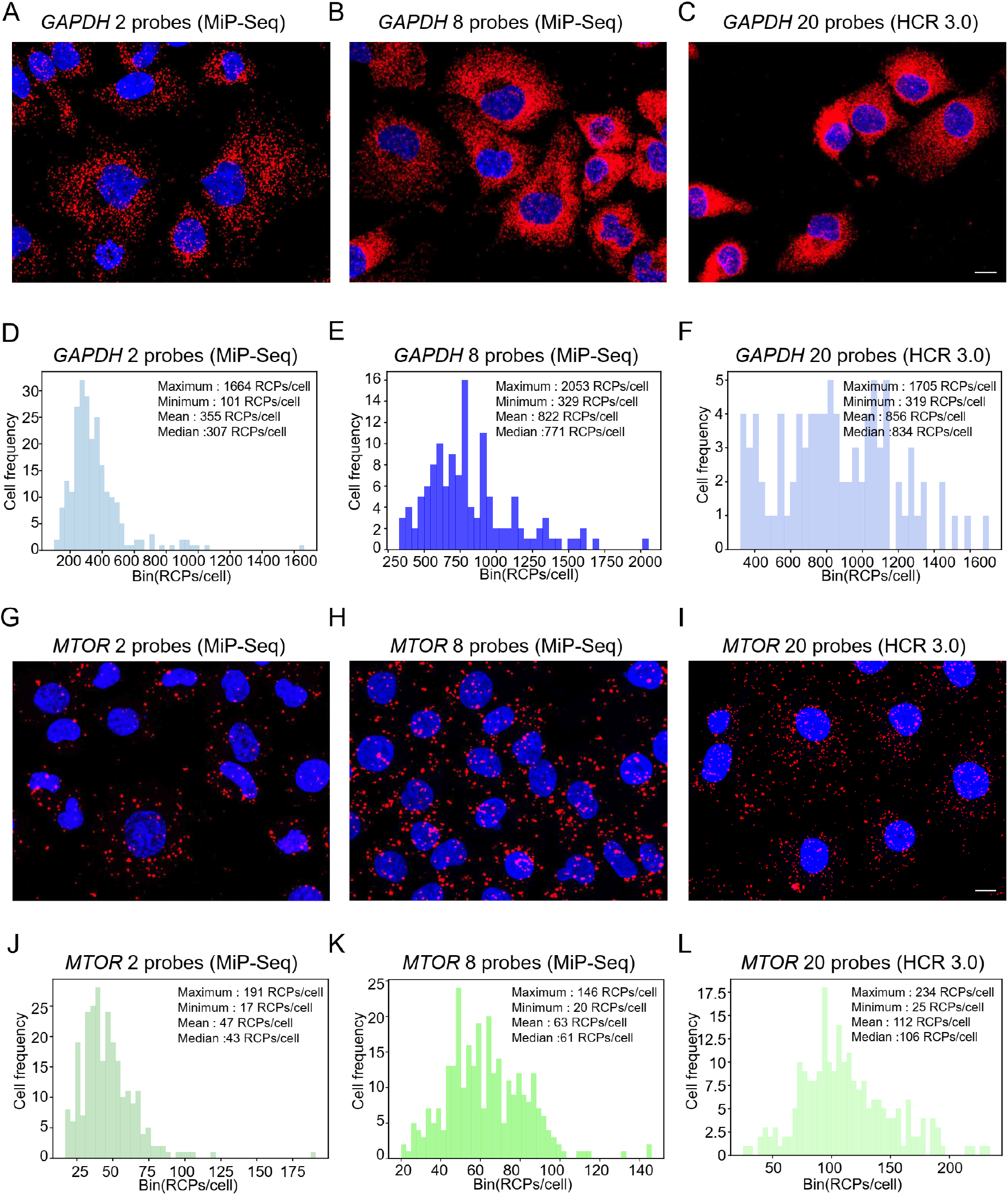
Comparison of detection efficiency between MiP-Seq and HCR 3.0. (**A-C**) Detection of *GAPDH* mRNA in HBMEC cells by MiP-Seq with 2 probe pairs (A), 8 probe pairs (B), and by HCR (3.0) with 20 probe pairs (C), respectively. (**D-F**) Diagram of signal distribution in single cells for A-C. (**G-I**) Detection of *MTOR* mRNA in HBMEC cells by MiP-Seq with 2 probe pairs (G), 8 probe pairs (H), and by HCR (3.0) with 20 probe pairs (I), respectively. (**J-L**) Diagram of signal distribution in single cells for G-I. Scale bars, 10 µm in A-C and G-I.

**Fig. S4.**
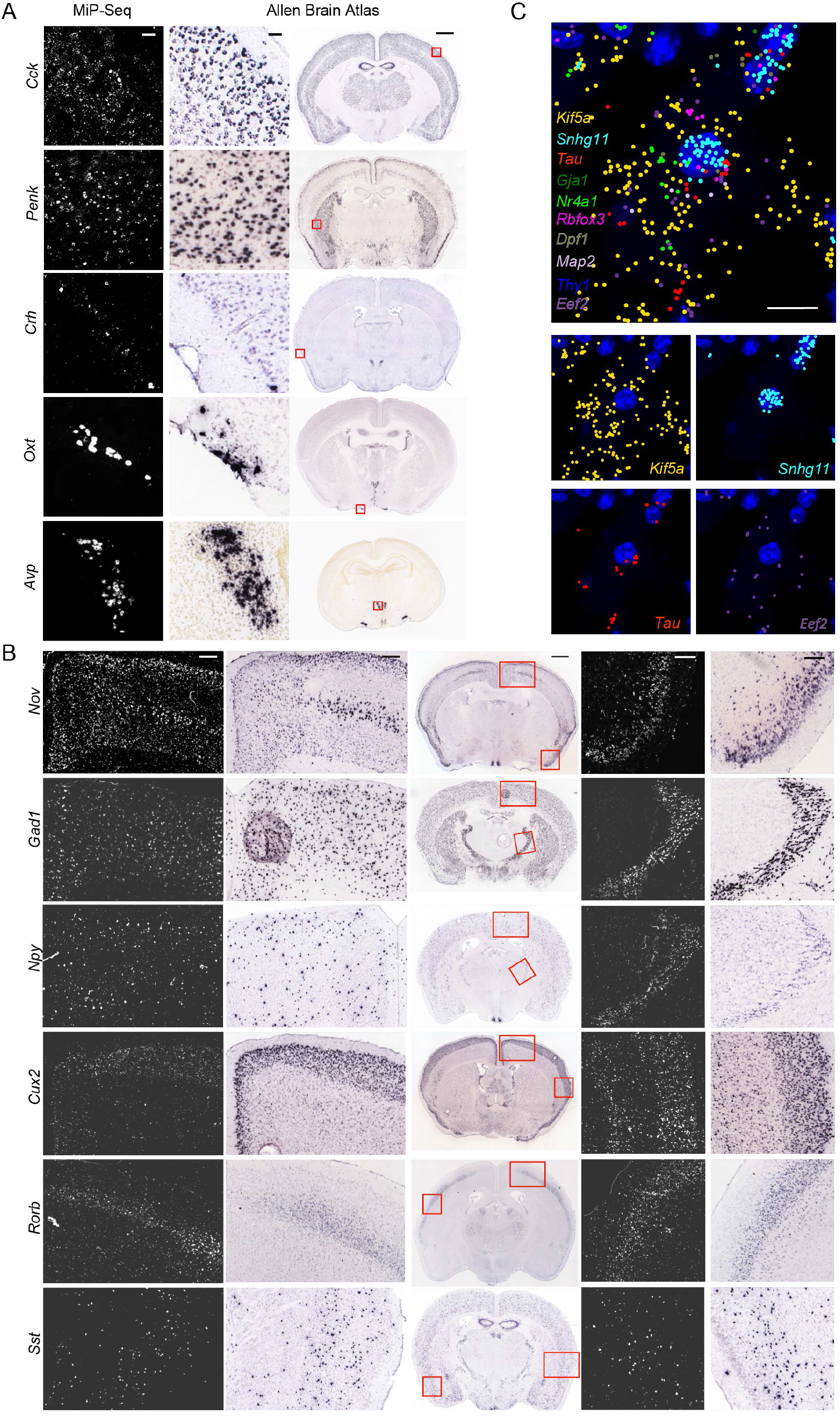
*In situ* detection of gene expression in brain slices via MiP-Seq. (**A-B**) The expression patterns of *Cck, Penk, Crh, Oxt, Avp, Nov, Gad1, Npy, Cux2, Rorb* and *Sst* genes detected by MiP-Seq were comparable to that by conventional *in situ* hybridization (from Allen Institute of Brain Science). (**C**) *In situ* co-detection of ten genes (*Kif5a, Snhg11, Tau, Gja1, Nr4a1, Rbfox3, Dpf1, Map2, Thy1*, and *Eef2*) in mice brain slices based on dual-barcode sequencing. Different pseudo-colors indicate the corresponding gene transcripts of ten genes, which display distinct subcellular localization (*Kif5a* in cytoplasm; *Snhg11* in nucleus). The subcellular localization of *Kif5a, Snhg11, Tau*, and *Eef2* genes transcripts are presented in the lower panels respectively. Scale bars, 50 µm in left two panels of A, 1000 µm in right panel of A and middle panel of B, 200 µm in left two panels and right two panels of B, 10 µm in C.

**Fig. S5.**
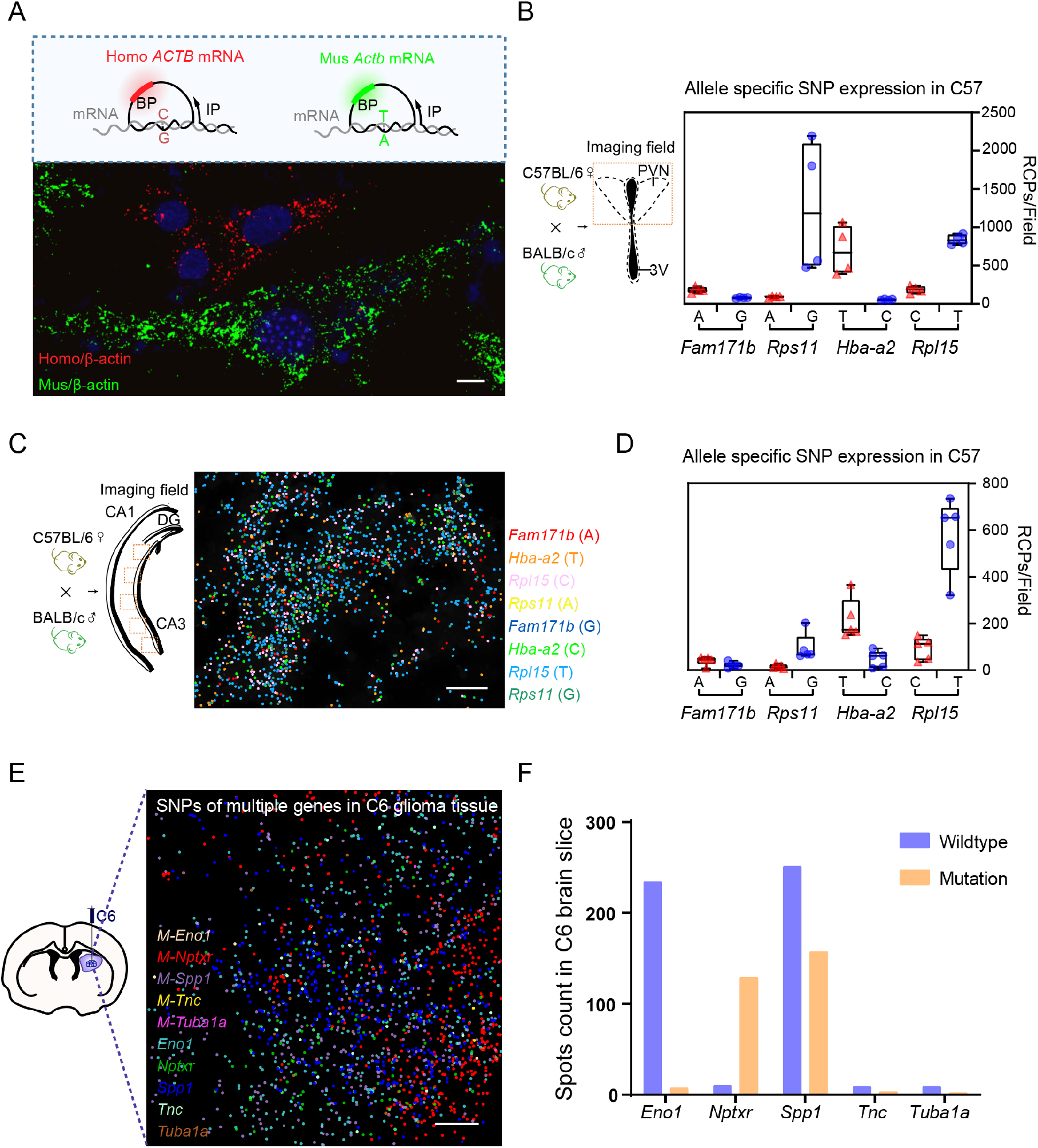
*In situ* detection of gene transcripts with single-base variance via MiP-Seq. (**A**) Detection of single-base variance in human and mouse β-actin gene transcripts in cocultured HBMEC (human origin) cells and Bend3 (mouse origin) cells. Red and green dot signals indicate human β-actin gene (*ACTB*) and mouse β-actin gene (*Actb*) mRNA sequence, respectively. (**B**) The box plot of the average gene expression level for *Fam171b, Rps11, Hba-a2*, and *Rpl115* alleles in PVN of the offspring mice from C57BL/6 male mice crossed with BALB/c female mice detected by MiP-Seq (based on the data from 4 fields of view). (**C**) *In situ* sequencing for allele-specific gene expression of *Fam171b, Rpl15, Hba-a2*, and *Rps11* in hippocampus of the offspring mice from C57BL/6 male mice crossed with BALB/c female mice. (**D**) The box plot of the average gene expression level for *Fam171b, Rps11, Hba-a2*, and *Rpl115* alleles in hippocampus of the offspring mice from C57BL/6 male mice crossed with BALB/c female mice detected by MiP-Seq (based on the data from 5 fields of view). (**E**) *In situ* co-detection of mutations for multiple genes (*Eno1, Nptxr, Spp1, Tnc*, and *Tuba1a*) in glioma area of rat brain from C6 glioma model. (**F**) The box plot shows the expression levels of wild-type and mutant variances of multiple genes in rat brain of C6 glioma model. Scale bars, 10 µm in A, 20 µm in C, 30 µm in E.

**Fig. S6.**
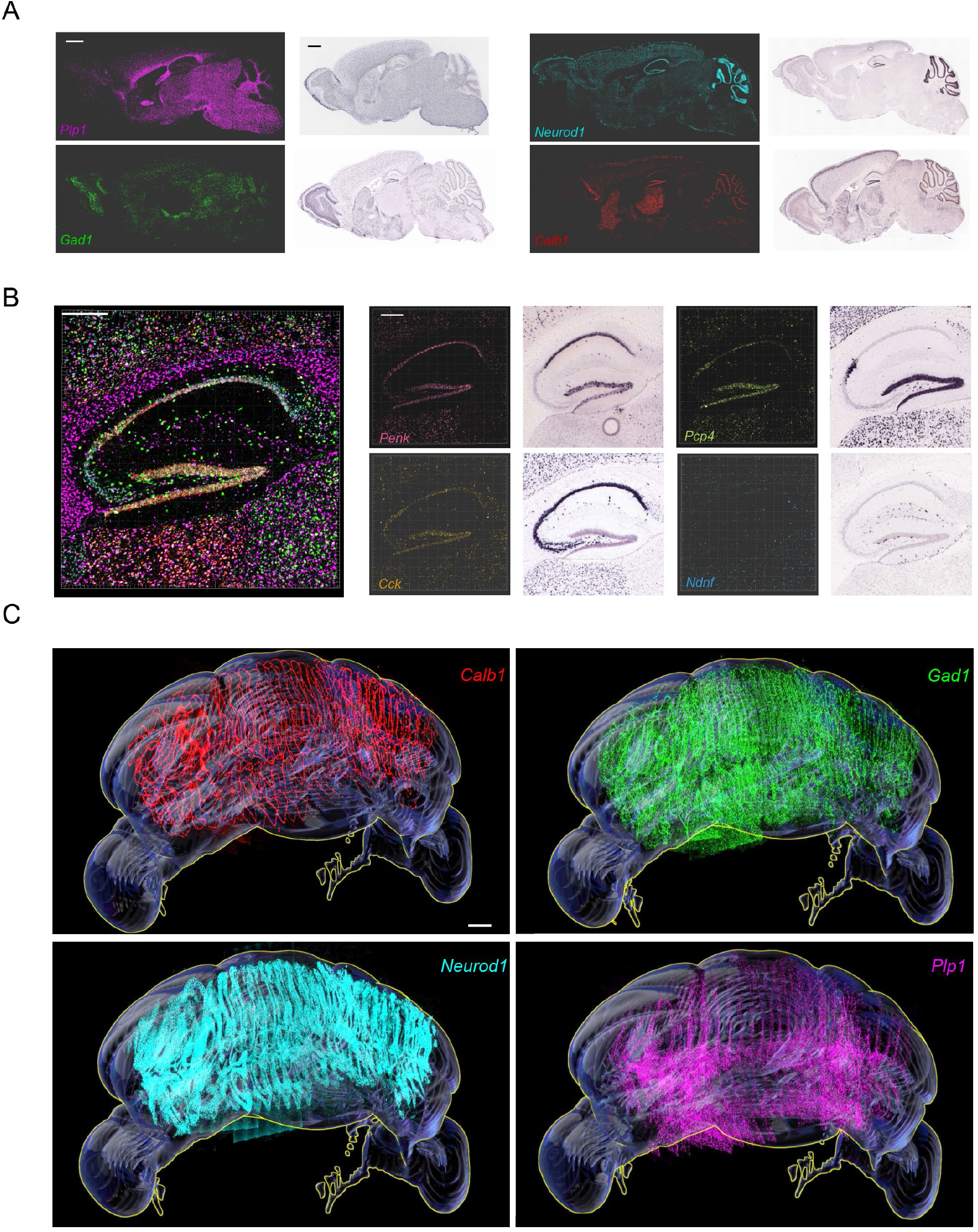
Spatial profiling of multiple genes in mice brain at a large-field of view and 3D reconstruction of the spatial profiling of specific marker genes in the whole cerebellum. (**A**) Expression of *Plp1, Neurod1, Gad1*, and *Calb1* in mice brain detected by MiP-Seq is consistent with conventional *in situ* hybridization results from Allen Institute of Brain Science. (**B**) The Left panel is the enlarged view of rectangle region in Fig.2A for ten genes codetection in hippocampus. The gene expression patterns in hippocampus detected by MiP-Seq are consistent with the results from Allen Institute of Brain Science (middle and right panels). (**C**) 3D view of *Calb1, Gad1, Neurod1*, and *Plp1* gene expression in the whole cerebellum are illustrated, respectively. Scale bars, 1000 µm in A, 300 µm in B, 500 µm in C.

**Fig. S7.**
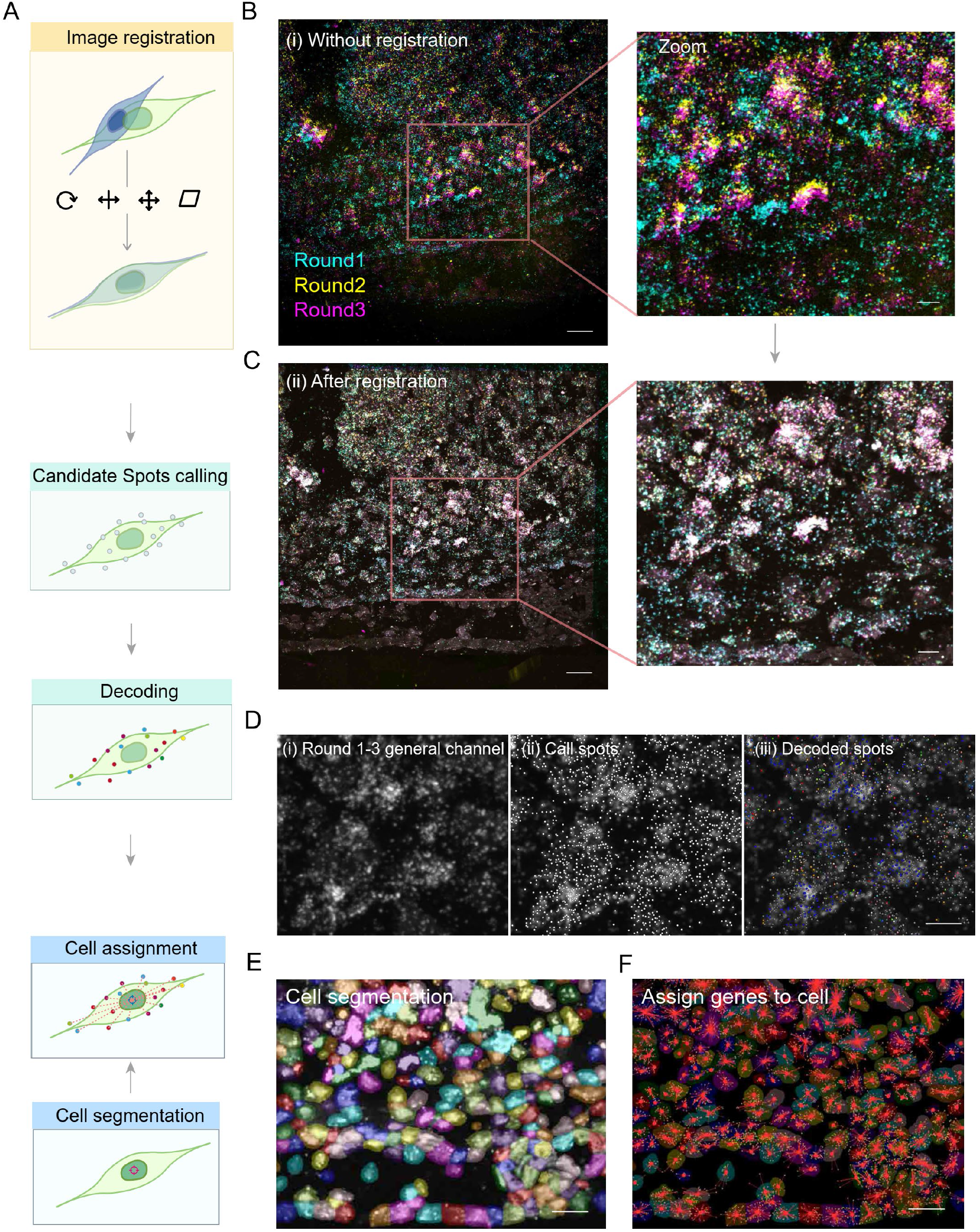
Image processing pipeline for spatial transcriptome analysis by MiP-Seq. (**A**) Diagram of image processing pipeline for spatial transcriptome analysis by MiP-Seq. The raw images from multiple rounds sequencing are first matched to the first-round image by affine transformations that allow rotation, translation, scaling, and shear transformations. The candidate points are selected according to the aligned images, and are decoded according to the intensity of each channel in each round at its location. The cell contour is calculated based on the aligned merged channel picture. Finally, the decoded gene points are allocated to neighboring cells according to the principle of proximity, and the expression profile corresponding to each cell is obtained. (**B**) The image (i) before registration. Three rounds of the channel merged images are represented in cyan, yellow, and magenta, respectively. (**C**) The image (ii) after registration. (**D**) Image and candidate spots in a local area. (i) Gray scale image of the combined channel images after alignment. (ii) Candidate spots are marked as white points on the image. (iii) Decoded spots are marked as points with different colors. (**E**) The cell mask is calculated from the cell morphology by cell pose. Different cells are indicated by different colors. (**F**) The results of gene assignment in this field of view. Different colored dots on the graph represent different genes. The red connecting line represents the gene to the corresponding cell. Scale bars, 30 µm in left image of B and C, 10 µm in right image of B and C, 10 µm in D, 30 µm in E and F.

**Fig. S8.**
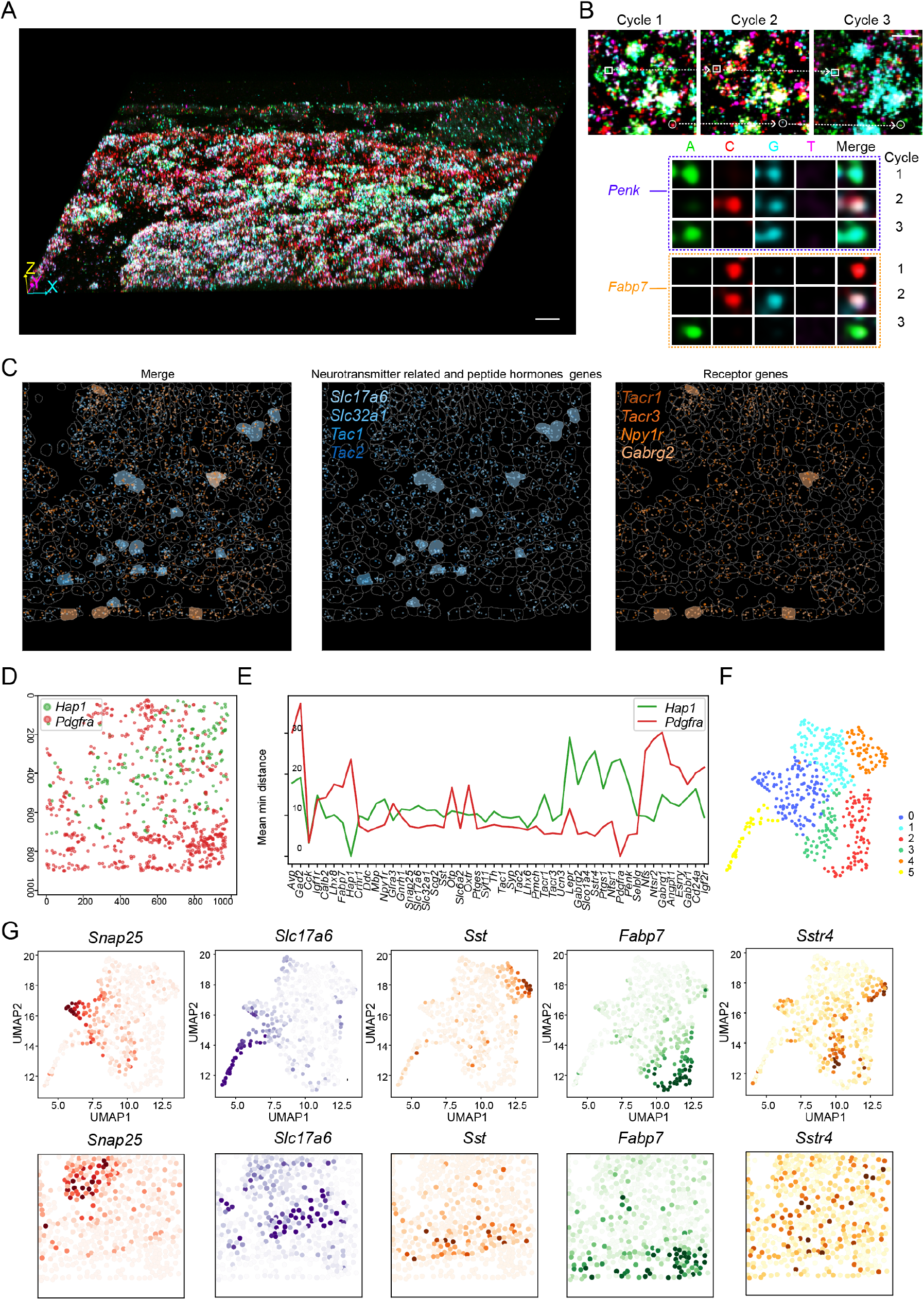
Spatial transcriptomic analysis of 100 genes in HDB by MiP-Seq and gene co-expression between cell clusters. (**A**) 3D projection image for *in situ* detection of 100 genes mRNAs in the first sequencing round by MiP-Seq in HDB. (**B**) The images of three round sequencing by MiP-Seq. The signals lined by the rectangle box and circle are further illustrated in the middle and lower panel respectively. (**C**) The spatial distribution of neurotransmitter-related and peptide hormone genes (blue) and their receptor genes (yellow). When the counts of the corresponding gene in each cell is greater than 10, the whole cell is marked by the color of the corresponding gene. (**D**) Spatial expression analysis of *Hap1* and *Pdgfra* genes in HDB. (**E**) The spatial distance of *Hap1* (green curve) and *Pdgfra* (red curve) gene to the top 50 expression genes. (**F**) Cells of HDB are clustered into 6 groups (0-5) based on the expression profile of single cells detected by MiP-Seq. (**G**) UMAP of featured genes expression in certain cell clusters and the corresponding spatial expression pattern in HDB. Scale bars, 50 µm in A, 10 µm in B.

**Fig. S9.**
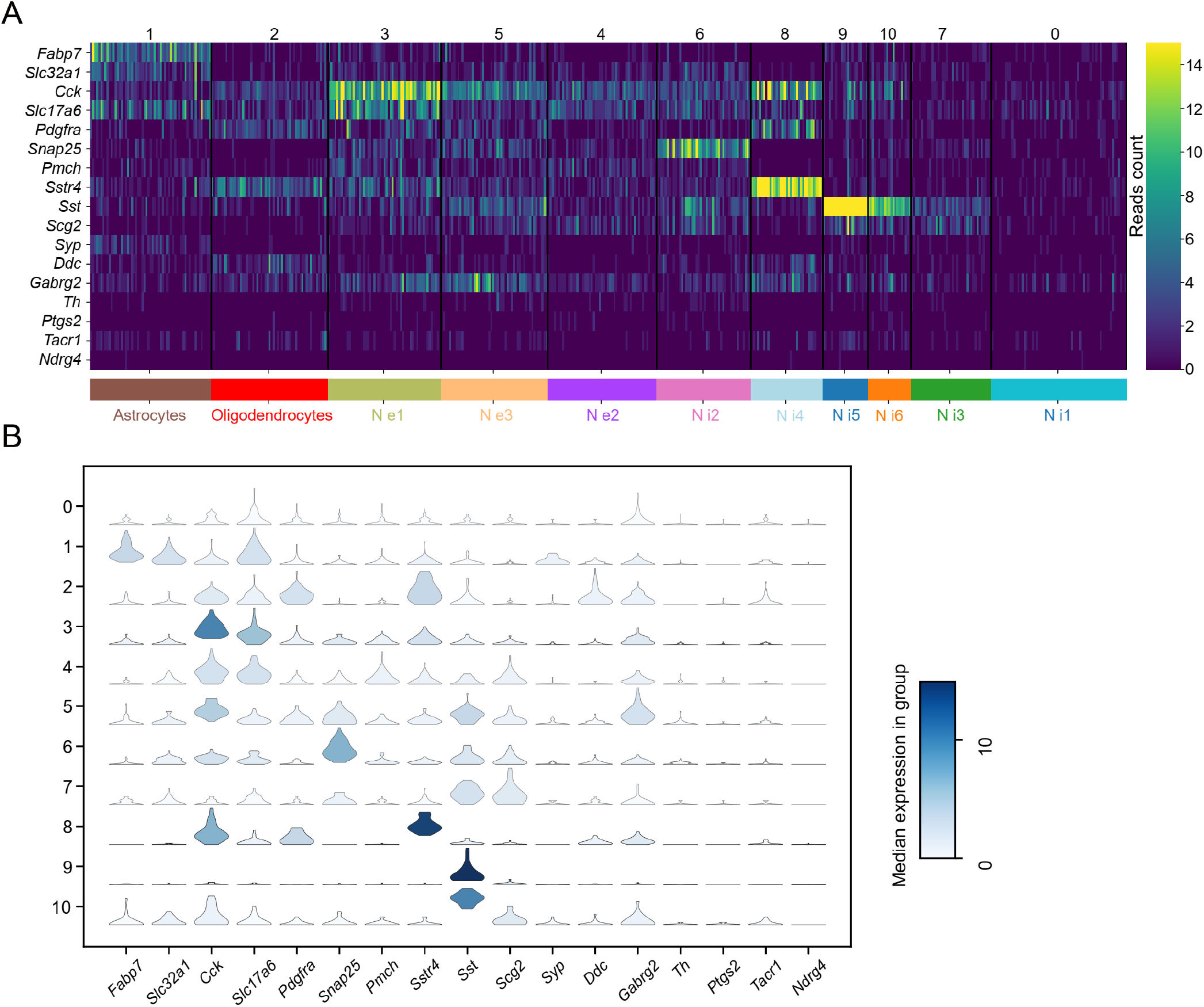
Single-cell clustering analysis. (**A**) Heatmap shows the expression level and identity of genes in 11 cell clusters including astrocytes, oligodendrocytes, three excitatory neuron subclasses, and six inhibitory neuron subclasses. (**B**) Violin plots of marker genes expression across 11 clusters.

**Fig. S10.**
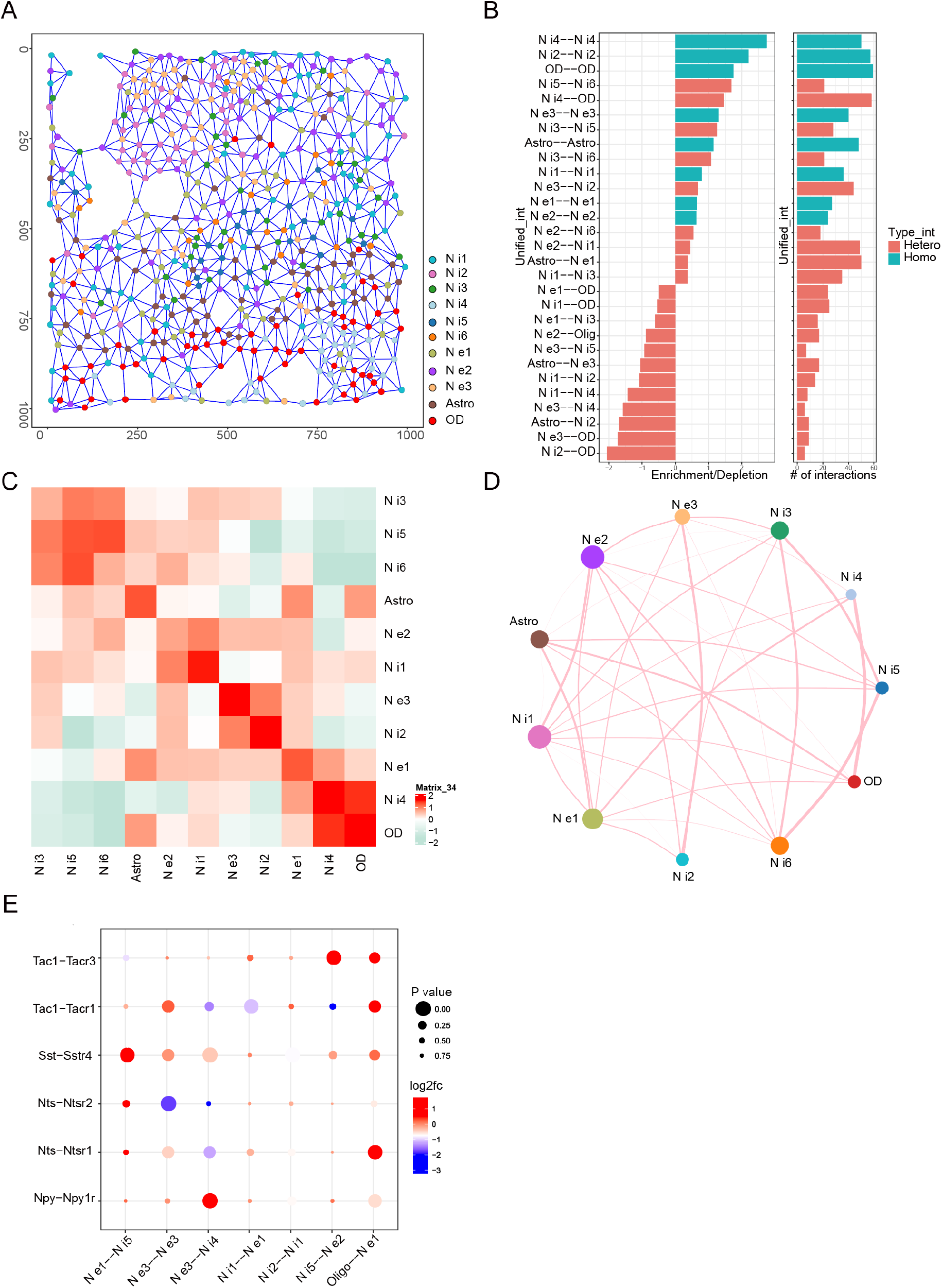
Analysis of spatial interaction relationship among cell clusters. (**A**) The spatial Delaunay network shows the physical distance between single cells. (**B**) The histogram (left) shows spatially proximal interactions between cell type pairs. On the x-axis, positive/negative values indicate enrichment/depletion (the observed frequency is higher/lower than the expected frequency). The histogram (right) shows the number of interactions. Blue indicates interaction within the same cell type, and red indicates interactions between different cell types. (**C**) Heatmap of spatial interaction among 11 clusters in HDB. (**D**) Cell-type interactions among different clusters. Lines connect to the cell population with spatial proximity enrichment. The thickness of the line indicates the strength of the interaction. Node size is positively correlated with the number of cell population pairs. (**E**) Spatial interaction of ligand and receptor among cells clusters. log2FC indicates the degree to which the expression level of the selected ligand-receptor pair is higher/lower than the expected level (log2(LR_expr/rand_expr)).

**Fig. S11.**
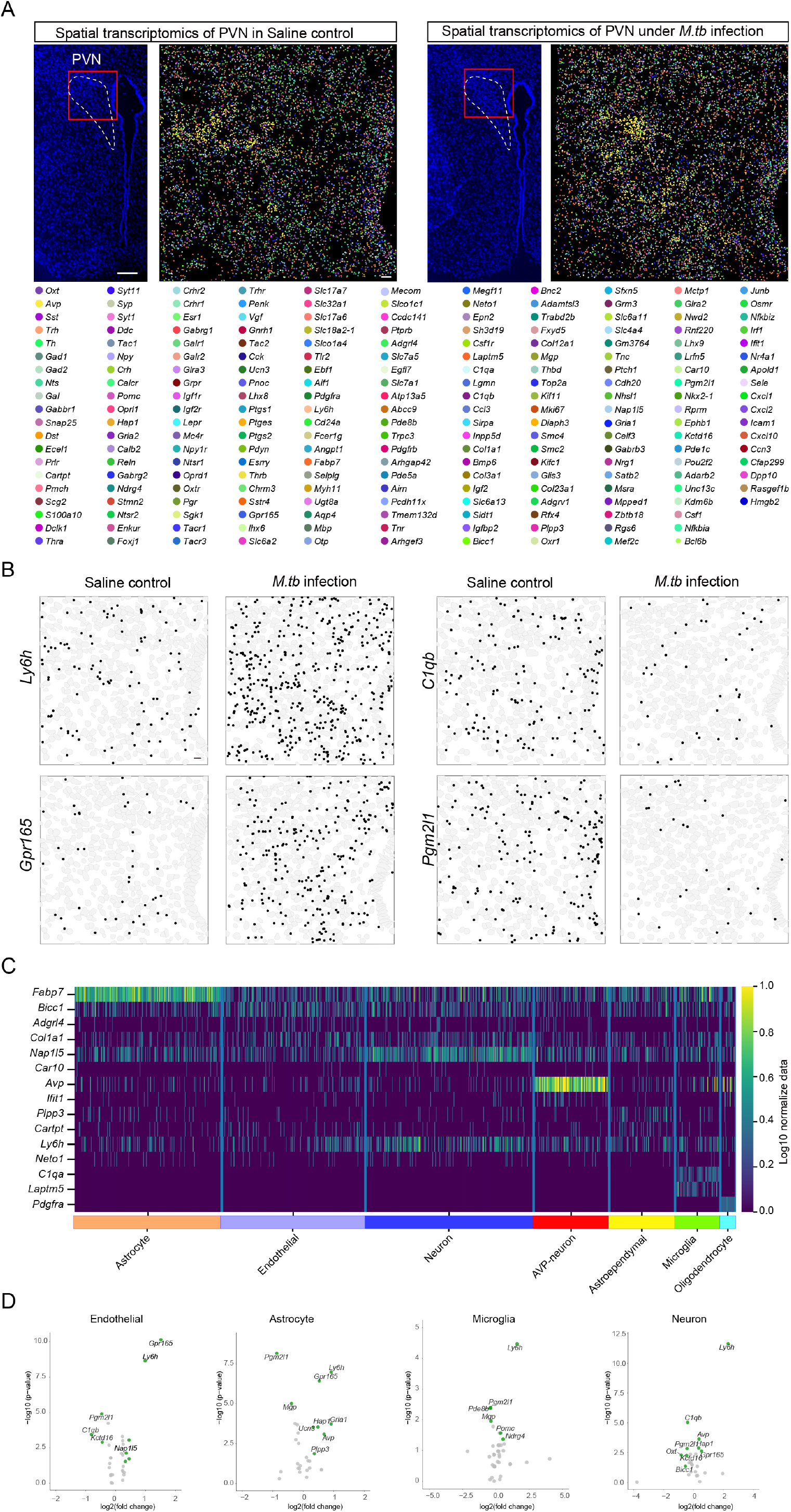
Spatial transcriptomic alteration of PVN upon *Mycobacterium tuberculosis* (*M*.*tb*) infection. **(A)** Detection of 217 genes including cell markers, hormones, receptors, as well as immune related genes in mice PVN upon *Mycobacterium tuberculosis* (*M*.*tb*) infection and saline control treatment by MiP-Seq. (**B**) Expression of *Ly6h* and *Gpr165* were up-regulated while *C1qb* and *Pgm2l1* were down-regulated in response to *M*.*tb* infection. (**C**) Heatmap of the expression of cell marker genes. (**D**) Differentially expressed genes in endothelial cells, astrocytes, microglia, and neurons. Genes with significantly changes (P value ≤0.05 and log(fold change) ≥0.3) were labeled in green. Scale bars, 100 µm in DAPI images of A, 10 µm in multiplexed genes detection image of A and in B.

**Fig. S12.**
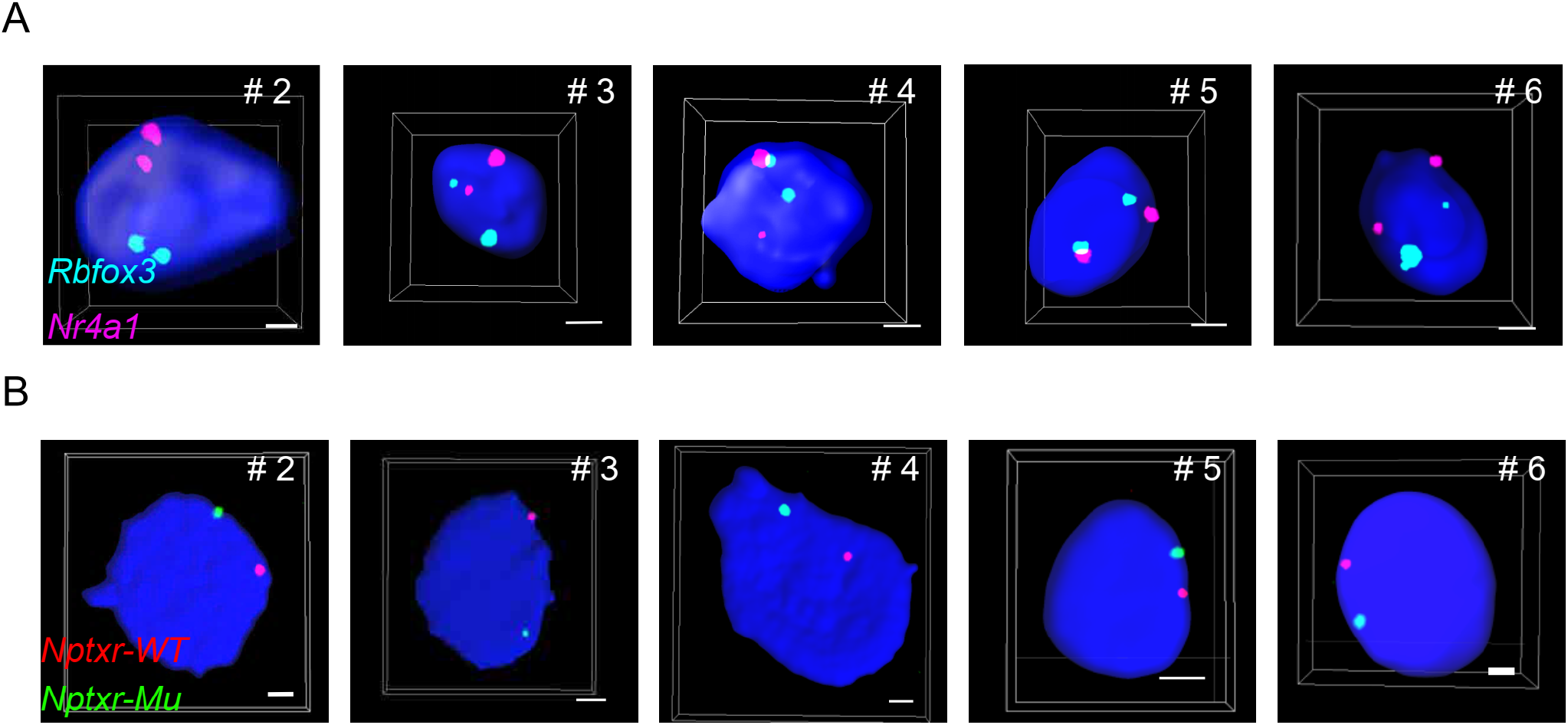
*In situ* detection of multiple genomic loci via MiP-Seq. (**A**) The remaining data for co-detection of *Rbfox3* and *Nr4a1* genes in mice tissue by MiP-Seq. Different cells are numbered. The data for #1 cell are shown in Fig.4B. (**B**) More images for *in situ* sequencing of *Nptxr* point mutation located in different chromosomes in C6 cell. Different cells are numbered. The data for #1 cell are shown in Fig.4C. Scale bars, 2 µm in A and B.

**Fig. S13.**
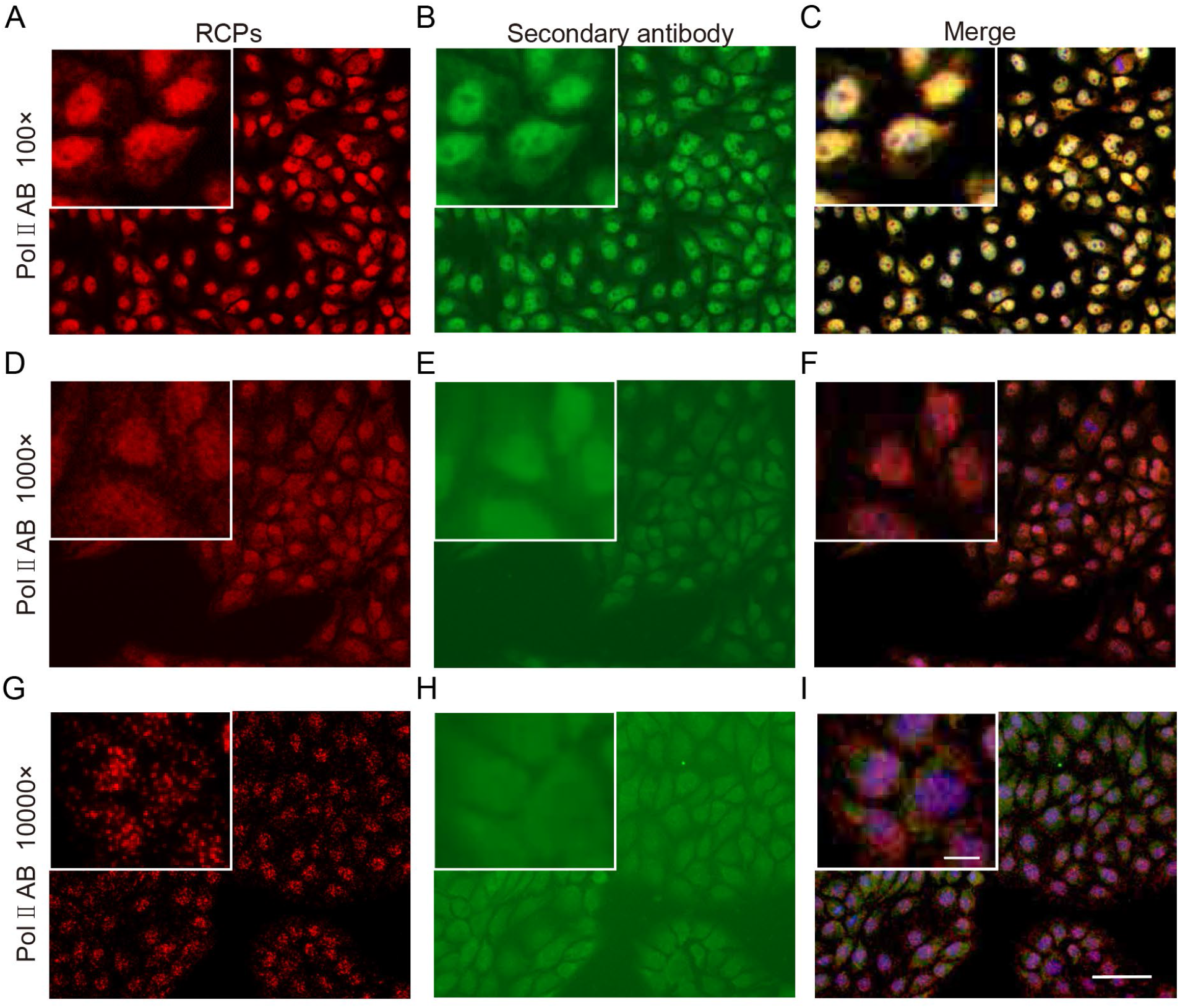
Higher sensitivity of protein detection via MiP-Seq in comparison to classic immunostaining. (**A, D, G**) Detection of RNA polymerase II (Pol II) protein in PK-15 cells via MiP-Seq at different dilution of primary antibody conjugated with nucleic acid. Strong signal for Pol II can be visualized, and clear dot signal can be observed under 10000 X dilution of primary antibody. (**B, E, H**) Detection of Pol II protein via classic immunostaining at different dilution of primary antibody. Positive signals were observed under 100 X dilution of primary antibody. After higher dilution, positive signal was hardly detected. (**C, F, I**) Merged images. Insets indicate higher magnification images. Scale bars, 50 µm for lower magnification, 10 µm for higher magnification.

**Fig. S14.**
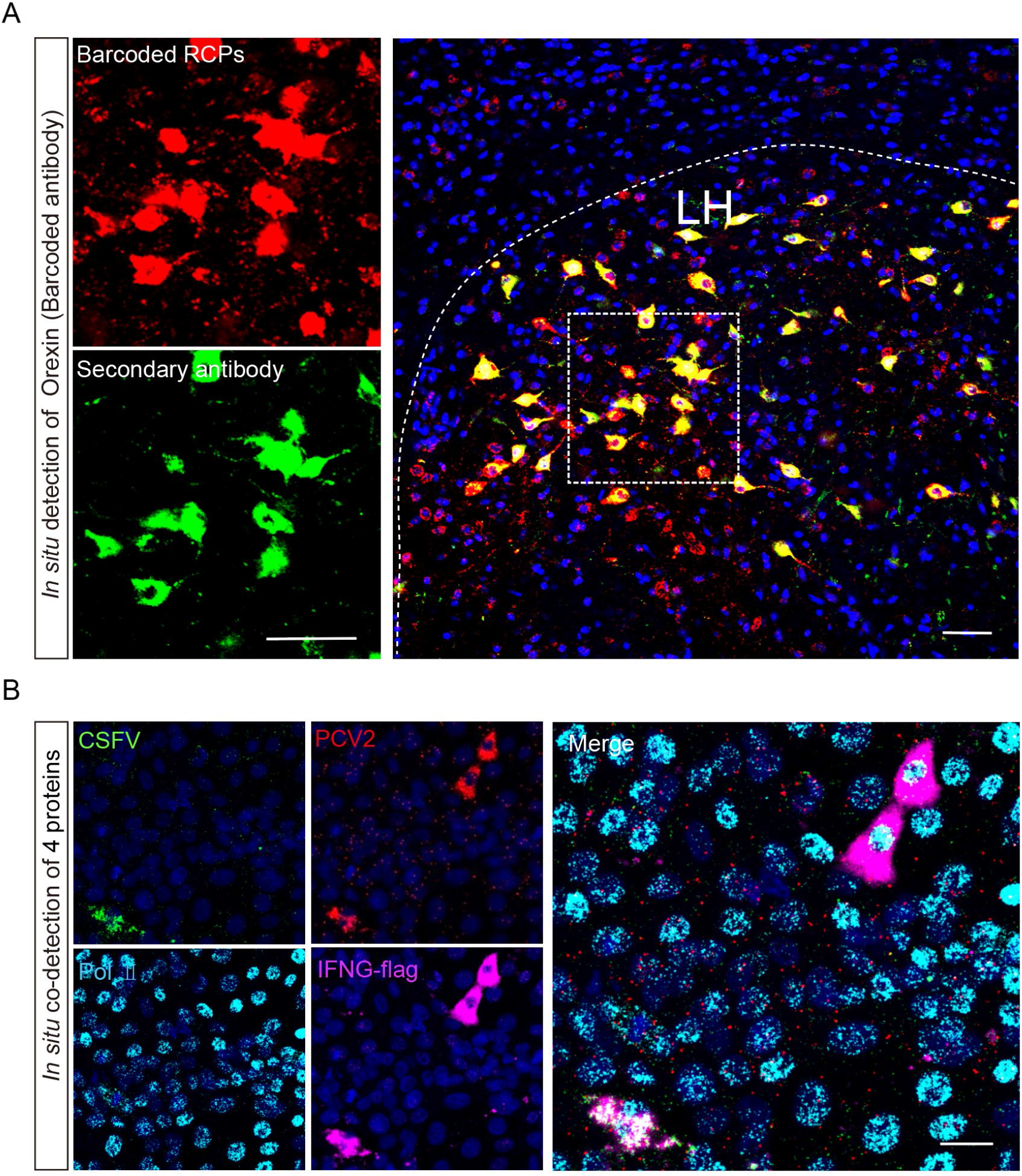
*In situ* detection of neuropeptide and multiple proteins via MiP-Seq. (**A**) Identical signals for neuropeptide orexin detected by MiP-Seq (Cy3 red fluorescence) and classic immunostaining (Alexa fluor 488 green fluorescence) were observed in lateral hypothalamus (LH) region of mice brain. For orexin detection by MiP-Seq, the targeting oligonucleotides was conjugated with secondary antibody. (**B**) *In situ* co-detection of four proteins (E2 of CSFV, Cap of PCV2, flag for IFNG, and Pol II) conjugated with distinct DNA oligonucleotides via MiP-Seq. Scale bar, 50 µm in A, 20 µm in B.

**Fig. S15.**
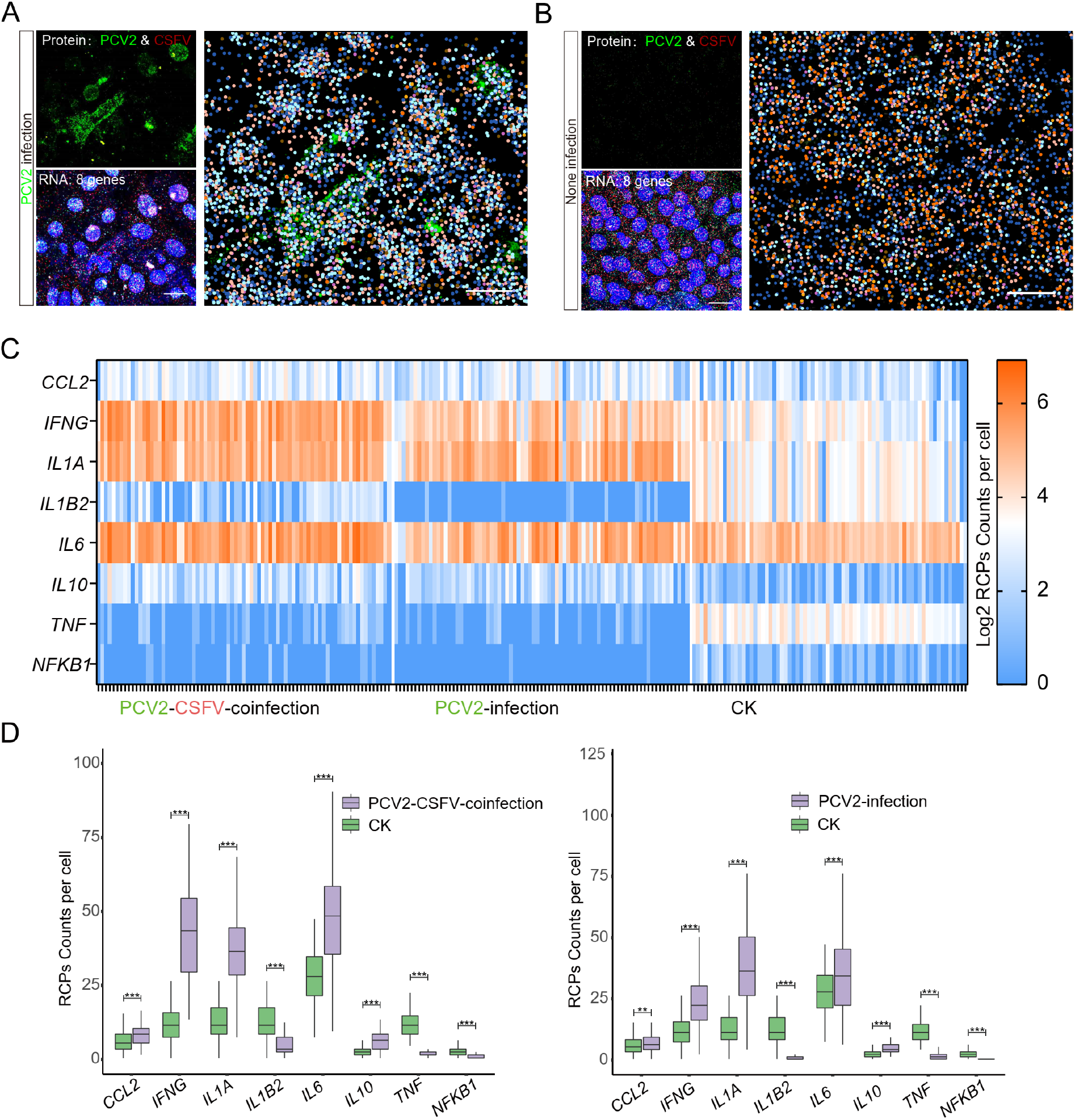
Simultaneous detection of cytokine/chemokine mRNAs and virus proteins via MiP-Seq. **(A)** Co-detection of mRNAs for eight cytokine/chemokine genes and virus proteins in PK-15 cells infected with PCV2. No CSFV protein but Cap protein for PCV2 was detected under PCV2 infection. Merged image was presented in higher magnification. **(B)** Co-detection of eight cytokine/chemokine mRNAs and two virus proteins in control PK-15 cells without virus infection. Merged image was presented in higher magnification. (**C**) Signals for eight cytokine/chemokine mRNAs detected by MiP-Seq were quantified by counting signal dot number in each PK-15 cell for PCV2/CSFV co-infection, PCV2 infection, and non-infection control, respectively. (**D**) Statistic analysis on differential expression level of eight cytokine/chemokine genes between PCV2/CSFV co-infection and non-infection control, or between PCV2 infection and non-infection control, respectively. Data was presented as mean ± SEM; **P<0.01, ***P<0.001. Scale bars, 20 µm in A and B.

**Fig. S16.**
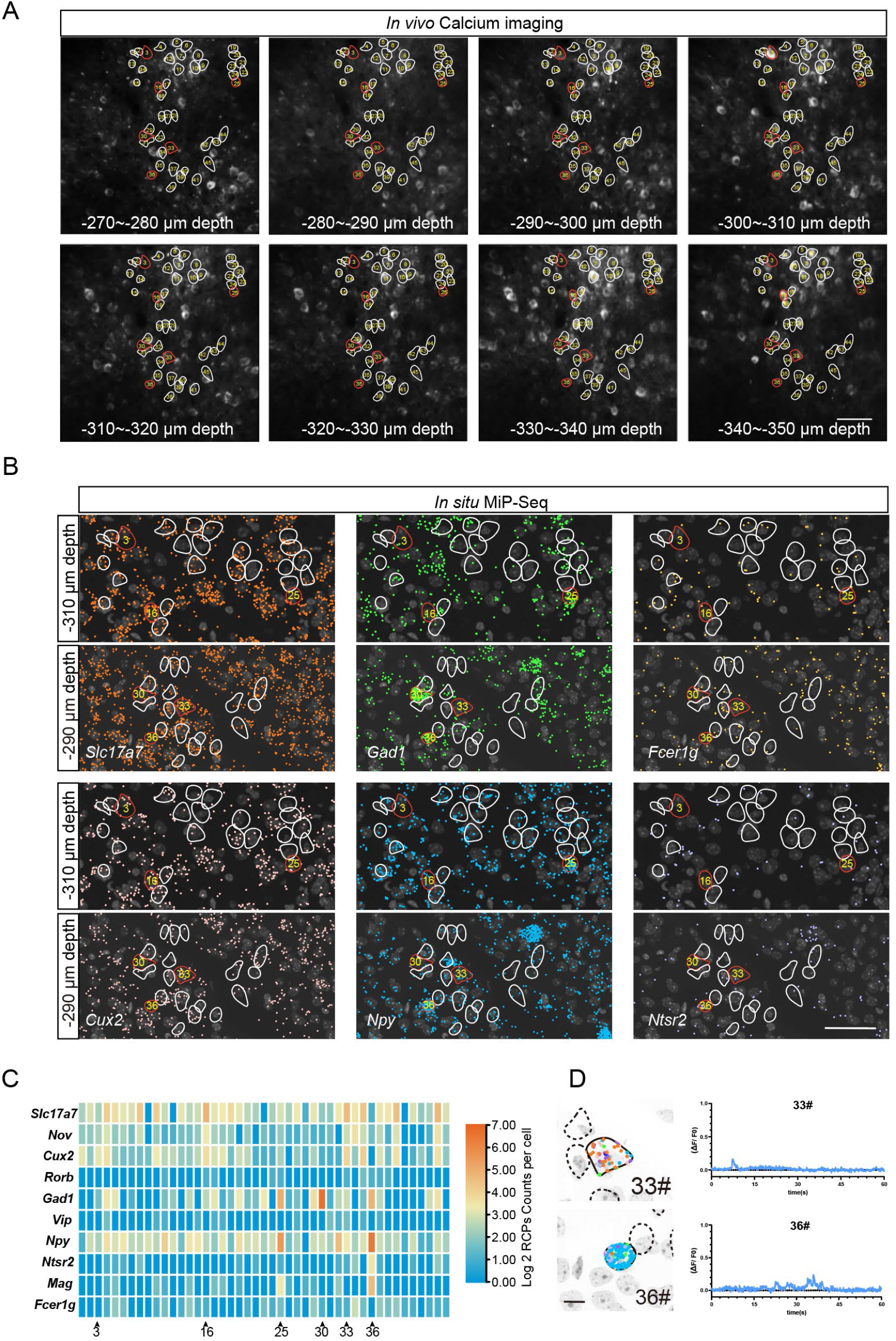
*In situ* analysis of the expression profiles of ten marker genes by MiP-Seq in calcium imaging area of mice V1 cortex. (**A**) Serial images of *in vivo* calcium imaging along 270∼350 µm depth of mice V1 cortex. (**B**) The expression profile of different marker genes including excitatory neuron marker genes (*Slc17a7* and *Cux2*), inhibitory neuron marker genes (*Gad1* and *Npy*), as well as microglia (*Fcer1g*) and astrocyte (*Ntsr2*) marker genes in calcium imaging area. (**C**) Quantification of signals for ten marker genes in 45 cells of calcium imaging regions by counting RCP number. (**D**) Correlation between gene expression profile and calcium signal response in cell 33# and 36#. Circles in A and B indicate cells for analysis of both transcriptomic profile and calcium dynamics, of which red circled cells were further provided with calcium dynamics curve. Scale bars, 50µm in A and B, 10 µm in D.

**Fig. S17.**
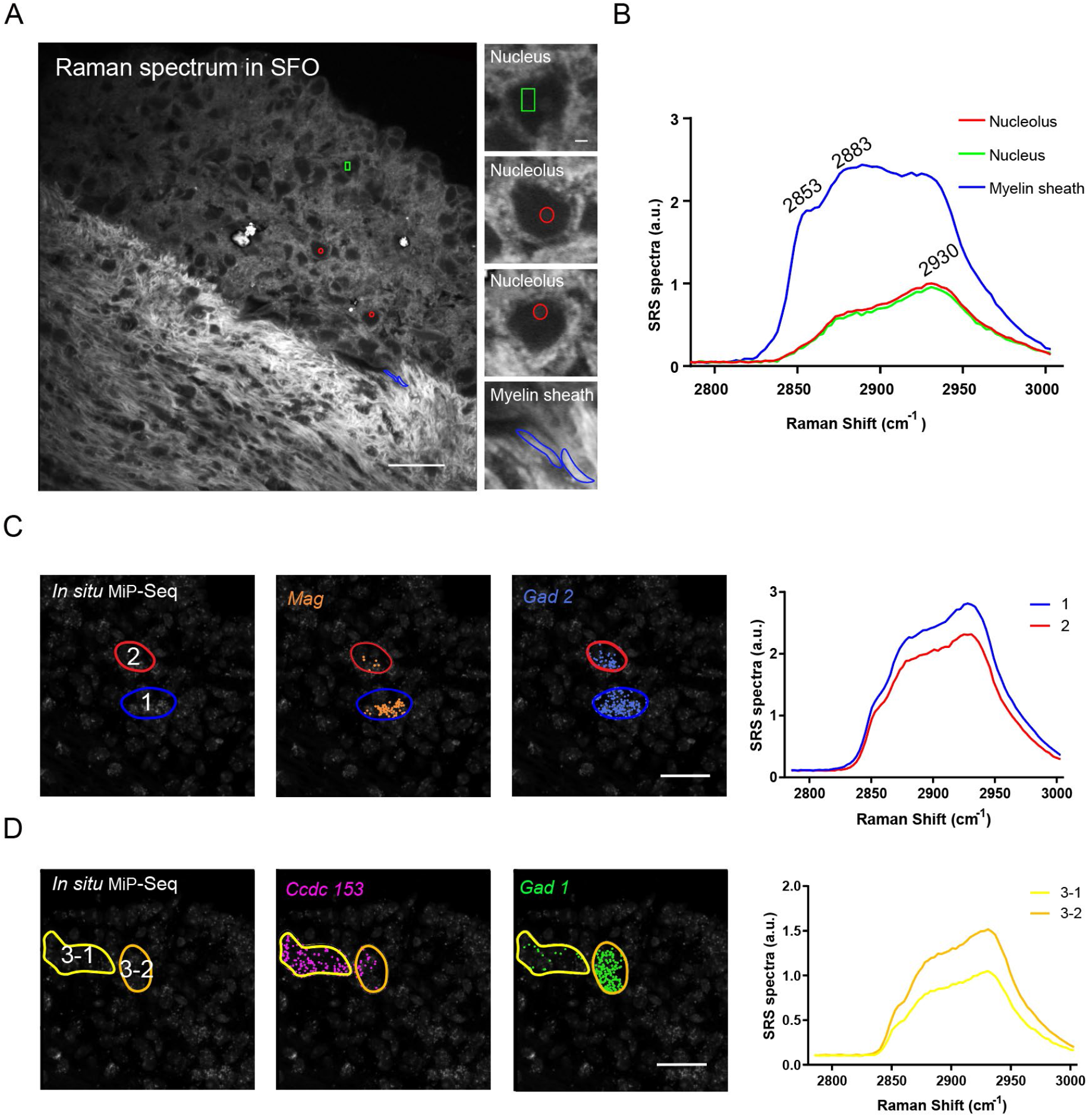
Integrated analysis of Raman spectroscopy imaging and marker genes expression profile detected by MiP-Seq. (**A**) The hyperspectral SRS of nuclei, nucleoli and fiber bundles image stack of cells in SFO. (**B**) The corresponding SRS spectra for a myelin sheath, nucleus, and nucleolus acquired in the relevant colored area indicated in (A). (**C**) Correlation between *Mag* and *Gad2* gene expression and certain chemical fingerprint regions (blue and red color region). (**D**) Correlation between *Ccdc153* and *Gad1* expression and certain chemical fingerprint regions (yellow and orange color regions). Scale bars, 30 µm in low magnification of A, 2 µm in higher magnification of A, 30 µm in C and D.

**Fig. S18.**
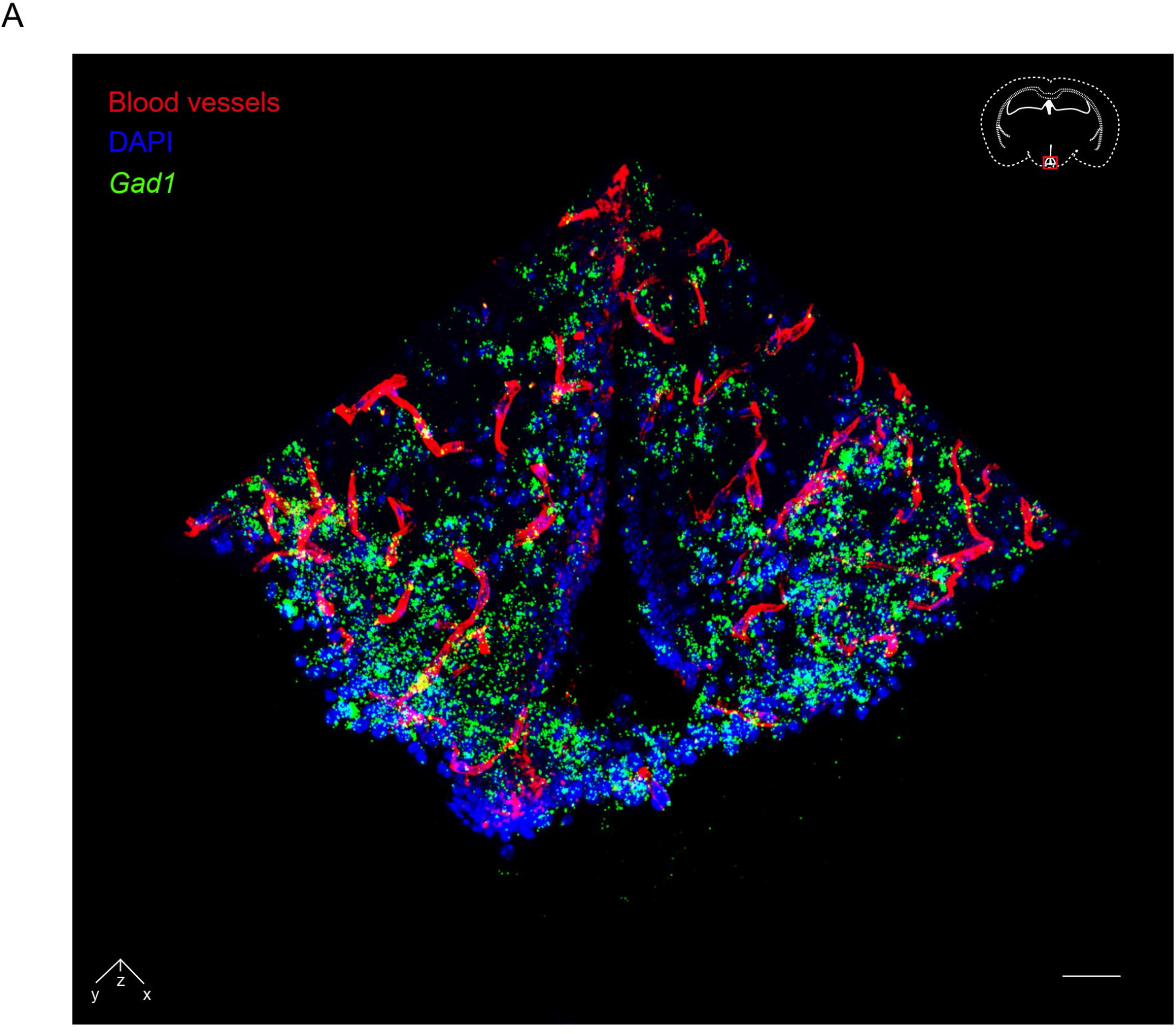
Simultaneous visualization of vasculature structure and gene expression. The blood vessels (red pseudocolor) stained by DyLight 488 Lycopersicon Esculentum Lectin and *Gad1* mRNA (green pseudocolor) detected by MiP-Seq in arcuate nuclei of mice brain were simultaneously visualized together with DAPI counterstaining. Scale bar, 50 µm.

**Fig. S19.**
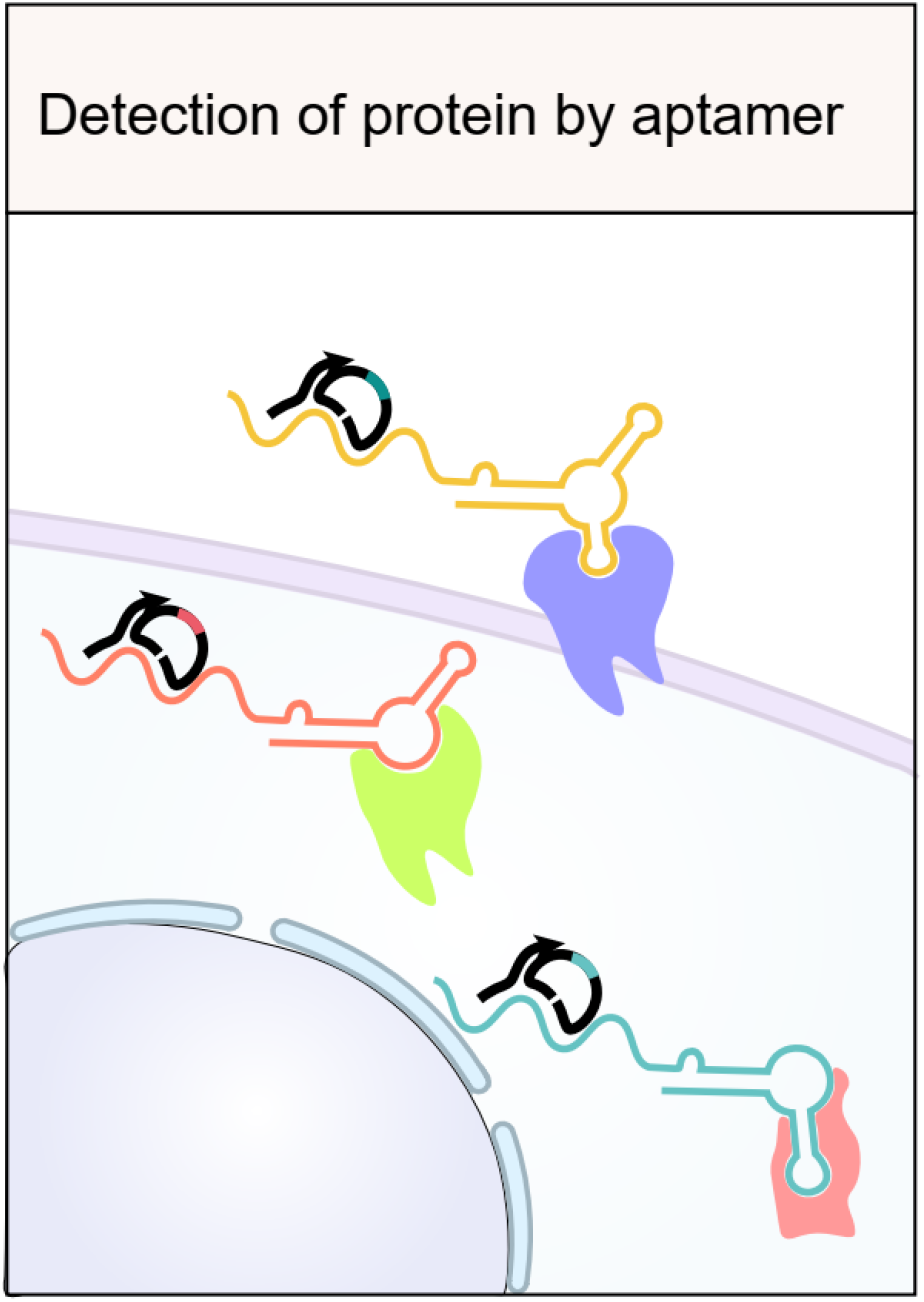
Aptamer based *in situ* detection of protein and biomolecule.

